# Computer vision-aided locomotor behavioral analysis identifies therapeutic motor signatures in a mouse model of Huntington’s disease

**DOI:** 10.64898/2026.09.03.749147

**Authors:** Lida Du, Zhenyu Wang, Qian Wu, Hongshuai Liu, Aadhya Bavkar, Yuan Zhou, Yukuan Shi, Lauren A Lim, Jiaxin Guo, Ziqi Qin, Christopher A Ross, Wenzhen Duan

## Abstract

Huntington’s disease (HD) is a neurodegenerative disorder characterized by progressive motor dysfunction. Traditional open-field tests quantify spontaneous locomotor parameters; however, fine mouse motor signatures, particularly disease stage-specific changes in HD motor symptoms and pharmacodynamic responses to therapeutic treatments. Here, we employed a computer vision-aided behavioral flow analysis designed to quantify fine, HD-relevant motor dysfunction in the zQ175DN HD mouse model, ranging from early HD-like motor signatures to well-defined motor deficits. Markerless pose estimation and Keypoint-MoSeq segmented standard top-view open-field recordings into recurrent behavioral syllables, which were then organized into higher-order clusters and transition networks. Disease stage-dependent changes in syllable occurrence, syllable duration, behavioral-state composition, and transition structure were identified. These analyses are not possible with traditional open-field assays. Syllable-duration features provided the strongest genotype discrimination, and HD-like motor features were also characterized by hub remodeling and transition-network disorganization. These features were integrated into an HD motor dysfunction (HDMD) score based on age- or HD progress-matched wild-type (WT) - standardized absolute deviations. The HDMD score distinguished HD mice from WT across multiple symptomatic stages and correlated with HD pathology and disease severity. Effect-size and power analyses suggested improved efficiency for detecting potential therapeutic effects. This framework requires only standard top-view recordings and may also support retrospective analysis of existing open-field video datasets. Overall, the HDMD framework provides a practical strategy for identifying fine motor changes in HD mice, aiding study design and preclinical efficacy assessment in HD drug development.

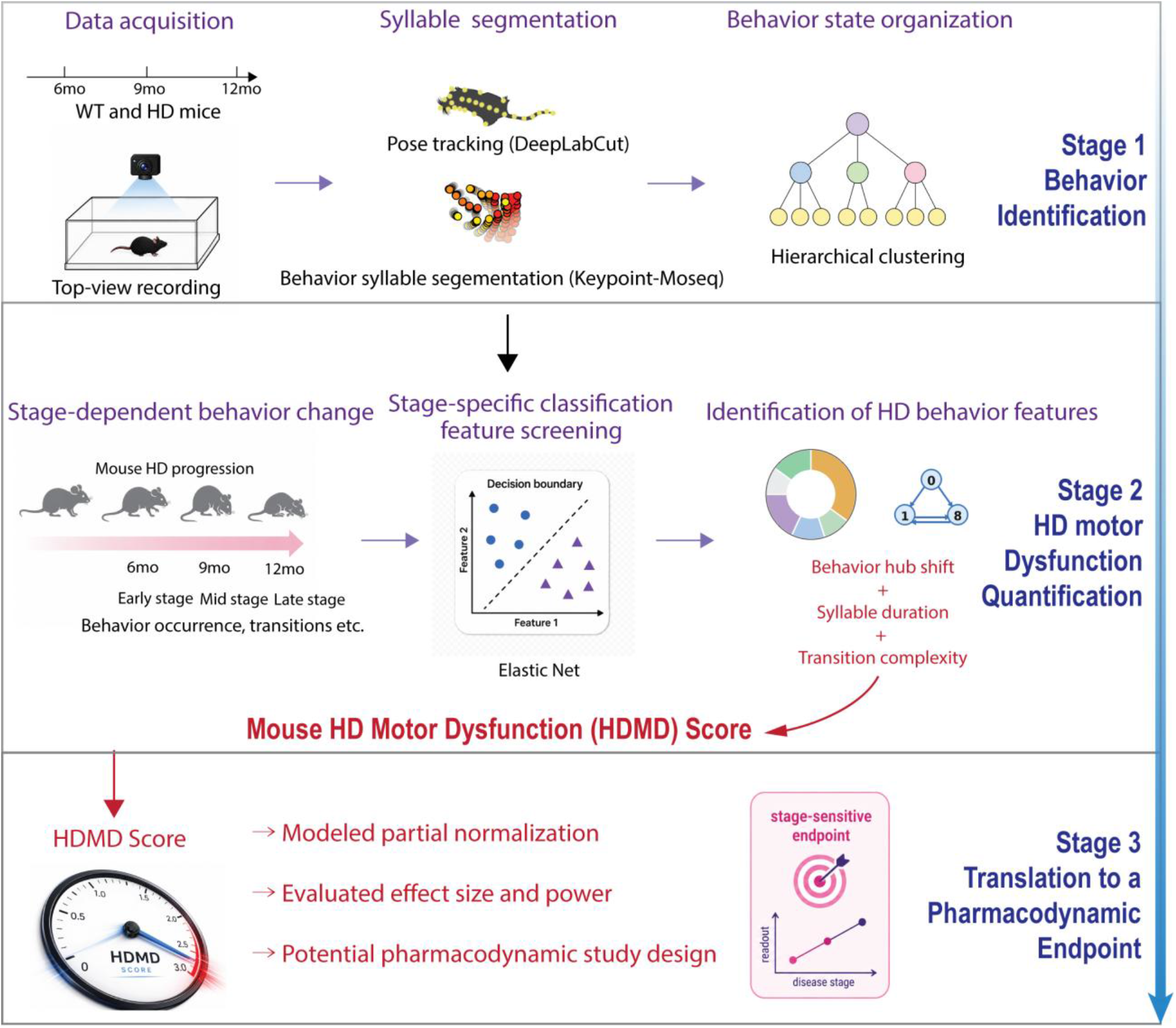

**Bullet Points:**

- Computer-aided behavioral analysis identifies stage-specific HD motor signatures in mice.
- The HDMD score provides clinically relevant motor assessment.
- HDMD improves power to detect therapeutic effects.
- Standard top-view videos may support retrospective HDMD analysis.

## Introduction

Huntington’s disease (HD) is an inherited neurodegenerative disorder caused by CAG-repeat expansion in the *HTT* gene and clinically characterized by progressive motor dysfunction, mental disturbance, and cognitive impairment ^1,2^. Corticostriatal disruption and striatal neuronal loss are key pathological features of HD progression^3–5^. Although HD mouse models reproduce key behavioral and neuropathological features^6,7^, preclinical drug development remains limited by the sensitivity and stage specificity of available locomotor endpoints^8^. A valuable endpoint measure should reproducibly detect HD-relevant subtle motor changes and is capable of showing pharmacodynamic responses to therapeutic treatment even before gross locomotor deficits become dominant.

The zQ175 DN knock-in mouse model is well suited for this purpose because mutant huntingtin is expressed in its endogenous genomic context and progressive behavioral and pathological abnormalities are characterized^6,9^. However, the robustness of motor phenotypes varies across disease stages and is affected by sex, body weight, and open-field test conditions^10–12^. Traditional behavior assays remain valuable, but they usually limit analysis to certain measures such as distance traveled or latency to cross or fall^12–14^. These measures provide limited information about how motor states are maintained and developed over disease stages. Thus, subtle treatment-responsive behavior changes may be overlooked. More sensitive and interpretable endpoints are therefore needed for stage-specific efficacy assessment in HD models.

Computer vision-aided video-based pose estimation tracks animal body parts without physical markers. When combined with data-driven behavioral segmentation, it resolves spontaneous movement at a finer temporal scale than conventional measures. Deep-learning pose-estimation tools, including DeepLabCut and SLEAP, enable body parts to be tracked in freely moving animals and enable quantitative analysis of naturalistic behavior^15,16^. Earlier supervised frameworks, such as JAABA, supported automated behavioral classification but relied on user-defined labels and were therefore sensitive to annotation bias^17^. More recent unsupervised methods, including B-SOiD, VAME, Motion Sequencing, and Keypoint-MoSeq, identify recurrent behavioral motifs directly from pose dynamics without predefined ethological categories^18–20^. In particular, DeepLabCut enables body landmarks to be tracked from standard laboratory videos, then Keypoint-MoSeq segments those tracked body trajectories into recurrent behavioral syllables^15,20^. Hierarchical clustering organizes these behavioral syllables into manually identifiable behavioral states based on their temporal sequence. This allows analysis of how often each state occurs, how long it is maintained, and how often to switch between behaviors^20–22^. Such approaches have revealed disease-related behavioral structure in other neurological disease models^23,24^.

Recent multi-camera 3D analysis in 8-week-old HD mice detected differences in a fine scale of motor abnormalities. However, that study was limited to a single prodromal HD stage ^25^. It remains unclear whether these behavioral signatures evolve during HD progression and serve as scalable preclinical endpoints. A scalable behavioral endpoint for future pharmacodynamic studies would ideally reflect clinically relevant motor symptoms across HD progression and integrate stage-dependent behavioral features. It would therefore be valuable to identify HD-defining behavioral features at each disease stage and develop them into a practical pharmacodynamic endpoint.

The compatibility of pose-based analysis with standard top-view videos also creates opportunities to extract more detailed behavioral features from existing open-field datasets^20,26^. However, their reliability across different datasets and recording conditions remains unclear.

In this study, computer vision-aided behavioral flow analysis was applied to zQ175DN model mice across early, intermediate, and developed HD. Behavioral syllables were organized into higher-order clusters and transition networks to define stage-specific motor signatures. Syllable-duration remodeling was most informative at earlier stages, whereas transition-network disruption became more prominent in later HD stages. Based on all these analyses, we integrated all these features into an HD motor dysfunction (HDMD) score from syllable-duration remodeling, behavioral hub shift, and transition complexity. We evaluated the HDMD statistical efficiency and potential application in preclinical pharmacological studies to support future HD pharmacological research.

## Results

### Computer vision-aided behavioral segmentation detected HD motor abnormalities beyond conventional open-field measures

One-hour top-view open-field recordings from 12-month-old zQ175DN HD mice and their litter-mate wild type (WT) were analyzed combining a series of computer vision-aided behavioral methods. Mouse pose was estimated with DeepLabCut^27^, and pose dynamics were segmented into recurrent behavioral syllables using Keypoint-MoSeq^28^. Conventional spatial measures, syllable occurrence, and syllable-transition structure were quantified from the same recordings (**Figure 1A**). Representative occupancy heatmaps showed genotype-dependent differences in spatial exploration with limited information on the fine-scale organization of motor behavior (**Figure 1B**).

**Figure 1.**
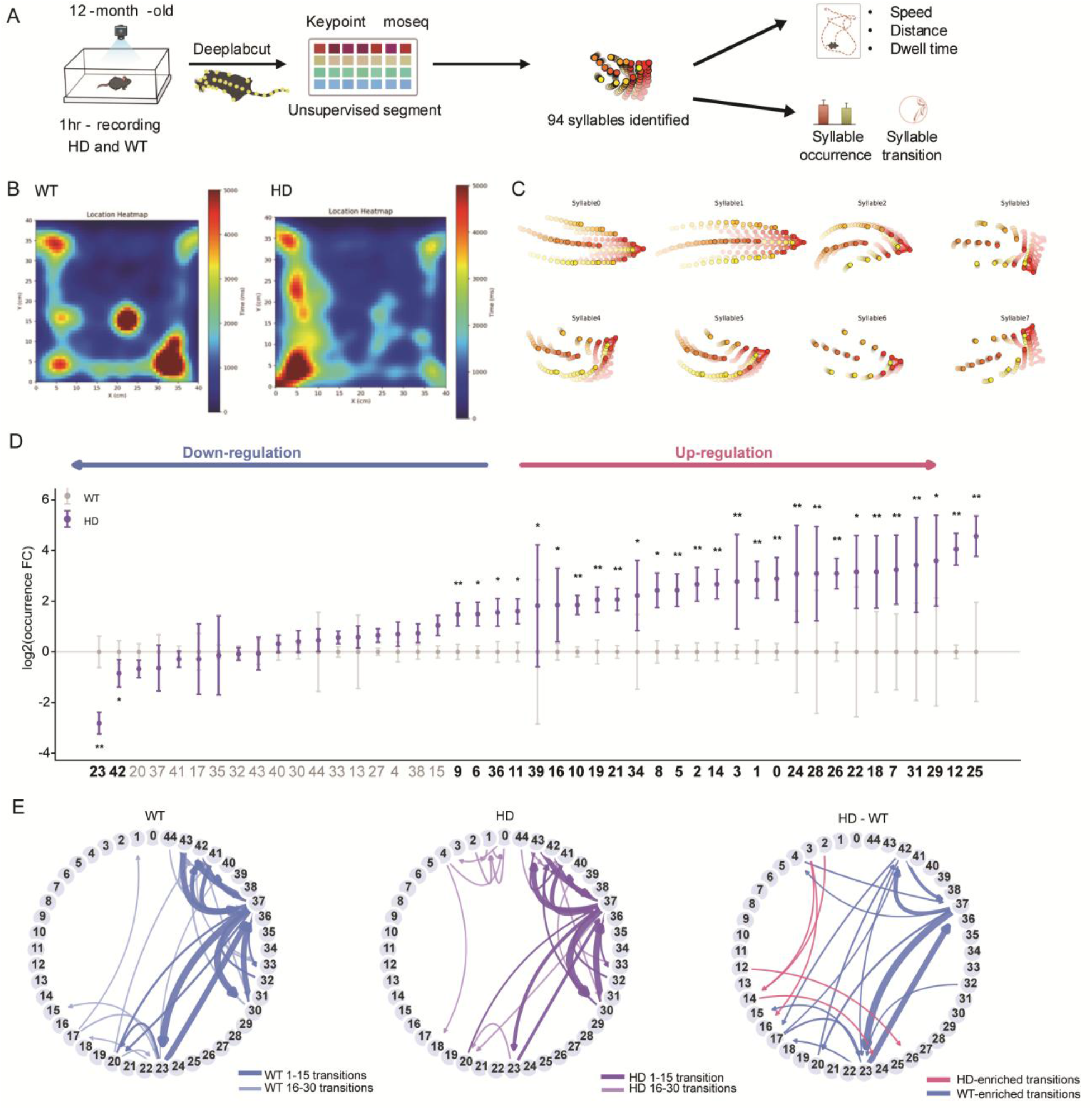
Computer vision-aided behavioral analysis identifies altered motor syllables and transition patterns in 12-month-old zQ175DN HD mice. **(A)** Workflow for open-field video acquisition and computational behavioral analysis. One-hour top-view recordings from freely moving WT and HD mice were processed using DeepLabCut-based pose estimation and Keypoint-MoSeq behavioral segmentation. **(B)** Representative spatial occupancy heatmaps from 12-month-old WT and HD mice. **(C)** Representative pose trajectories illustrating recurrent behavioral syllables identified by Keypoint-MoSeq. **(D)** Differential occurrence of behavioral syllables in 12-month-old HD mice relative to WT mice, shown as log₂ fold change. **(E)** Syllable-transition networks in WT and HD mice and the HD–WT differential network. Nodes represent behavioral syllables, and edges represent transition relationships between syllables. Data are shown as mean ± SEM; each point represents one mouse. Statistical analysis was performed using multiple t-tests where appropriate. For syllable-level comparisons, statistical analysis was performed between 12-month-old WT and HD mice. *p < 0.05, **p < 0.01, ***p < 0.001.

Keypoint-MoSeq initially identified 94 recurrent syllables in the discovery dataset. After exclusion of rarely expressed syllables, 45 syllables that occurred more than 96% frequency in total behavior were retained for downstream analyses. Representative pose trajectories illustrate the distinct movement patterns captured by individual syllables (**Figure 1C**), and trajectories of the 42 most frequently expressed retained syllables are provided in **Figure S1**. Differential occurrence analysis revealed bidirectional changes in 12-month-old HD mice, with two syllables occurring less frequently and multiple syllables occurring more frequently than in WT mice (**Figure 1D**). Thus, the HD phenotype was not characterized by uniform suppression of spontaneous behavior but by selective redistribution across motor syllables.

Syllable-transition networks further showed that genotype-associated abnormalities extended beyond the occurrence of individual states. Distinct transition patterns were observed in WT and HD mice, and the HD-minus-WT differential network identified both HD-enriched and WT-enriched transition routes (**Figure 1E**). Together, these findings demonstrate that computer vision-aided behavioral segmentation resolves HD-associated alterations in both the expression and sequential organization of spontaneous motor states that are not apparent from spatial occupancy patterns alone.

### Temporal hierarchical clustering resolved interpretable HD-associated behavioral states

Hierarchical clustering was performed according to the syllable temporal organization within the behavioral sequence to higher-dimensional behavioral states^29^. Syllables with similar sequential occurrence patterns were grouped into nine behavioral clusters, which were annotated from their representative pose trajectories as head turning, body turning, turning locomotion, slow rotation, excessive limb movement, segmented locomotion, fast rotation, sedation, and upright activity (**Figure 2A**).

**Figure 2.**
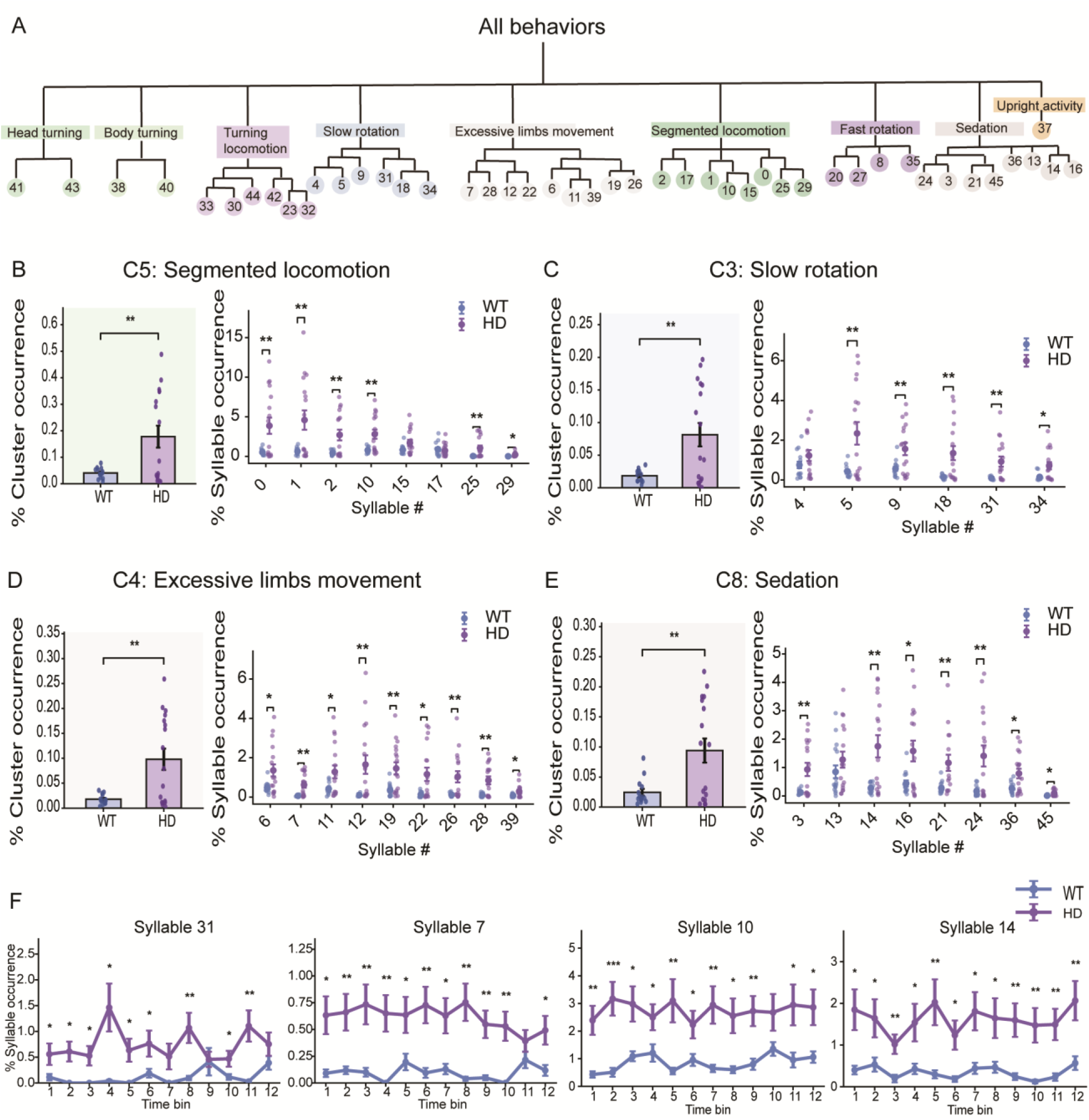
Hierarchical clustering of locomotor behavior reveals HD-associated movement patterns. **(A)** Hierarchical clustering of Keypoint-MoSeq syllables according to their temporal organization within the behavioral sequence, showing major behavioral clusters in the 12-month discovery dataset. **(B–E)** Occurrence of HD-associated behavioral clusters and their component syllables in 12-month-old WT and HD mice. HD mice showed increased occurrence of **(B)** segmented locomotion, **(C)** slow rotation, **(D)** excessive limb movement, and **(E)** sedation-related syllables**. (F)** Temporal occurrence patterns of representative HD-associated syllables across recording bins in 12-month-old WT and HD mice. Data are presented as mean ± SEM. Each point represents one mouse. Statistical significance was assessed by multiple t-tests. *P < 0.05, **P < 0.01, ***P < 0.001.

Cluster-level analysis identified several behavioral states that were enriched in 12-month-old HD mice (**Figure S2**). The occurrence of segmented locomotion was increased, with significant increases observed in multiple component syllables within this cluster (**Figure 2B**). The frequency of slow rotation was increased, accompanied by enrichment of several constituent syllables (**Figure 2C**). Increased cluster occurrence was also detected for excessive limb movement and sedation, with multiple syllables contributing to each effect (**Figure 2D,E**). These findings indicate that the cluster-level abnormalities were distributed across several related motor syllables rather than being driven by a single motif.

Temporal-bin analysis further showed that representative HD-enriched syllables remained elevated across multiple intervals of the one-hour recording session (**Figure 2F**). Representative videos illustrate segmented locomotion in WT and HD mice (**Videos S1 and S2**, respectively) and slow rotation in WT and HD mice (**Videos S3 and S4**, respectively). Together, these results demonstrate that temporal hierarchical clustering converted the fine-scale syllable vocabulary into interpretable behavioral states and revealed sustained enrichment of segmented locomotion, slow rotation, excessive limb movement, and low-activity-related behavior in developed HD.

### Altered locomotor motif structure and syllable-transition organization are identified in HD mice

Individual behaviors clusters are composed of single or multiple syllables; we further examined the syllable organization structuring a higher-dimensional locomotor behavior. For example, a complete body-rotation sequence can be combined by multiple Keypoint-MoSeq syllables (**Figure 3A**). This analysis framework allowed behaviors to be interpreted not as isolated syllables, but as structured sequences of pose-defined motor elements.

**Figure 3.**
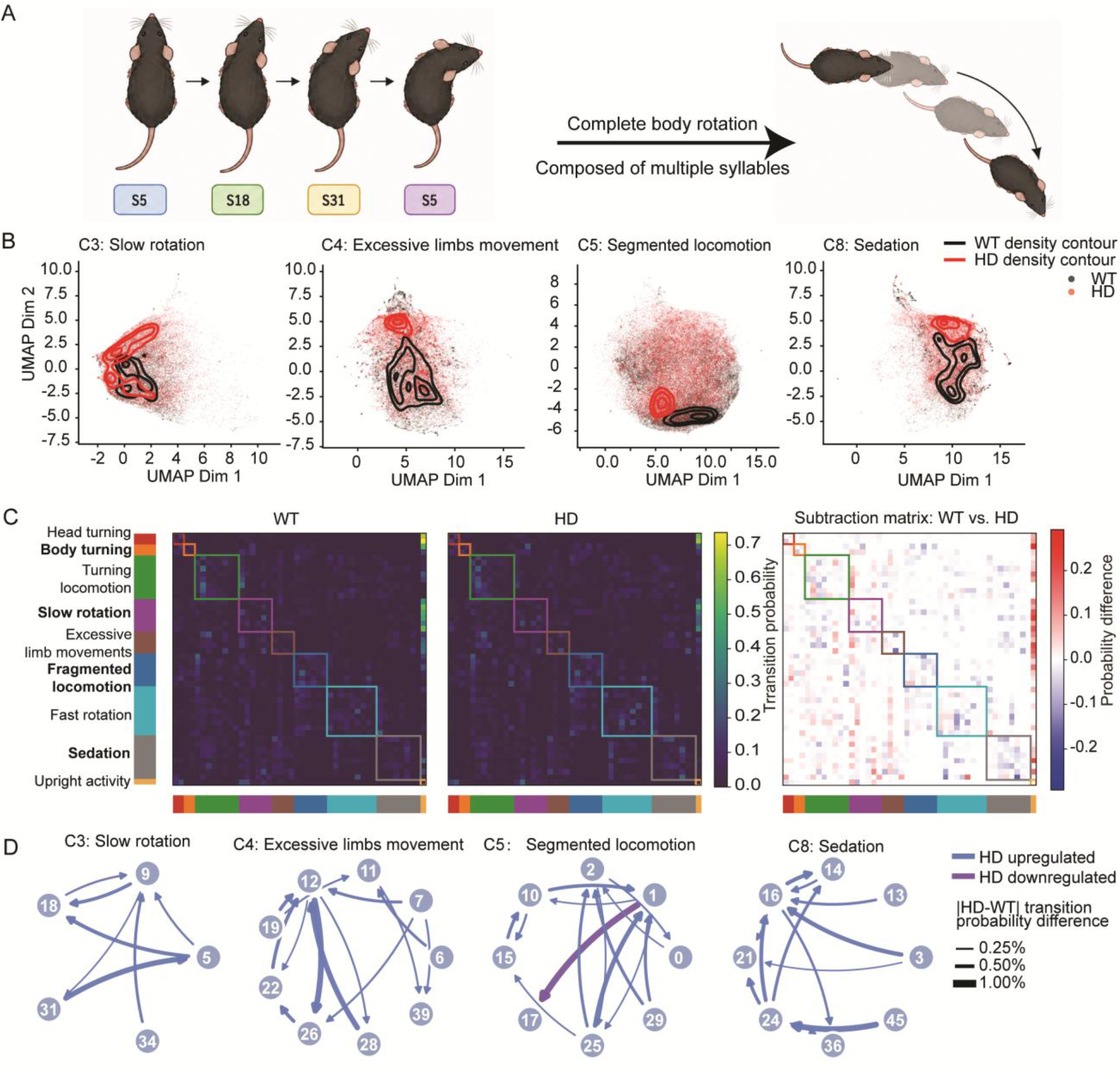
HD mice exhibit altered locomotor structure and motif transition patterns. **(A)** Schematic example showing how individual Keypoint-MoSeq syllables can combine into a higher-order locomotor motif, illustrated by a complete body-rotation sequence composed of multiple syllables. **(B)** UMAP visualization of pose-defined syllable distributions within representative behavioral clusters, including slow rotation, excessive limb movement, segmented locomotion, and sedation. Density contours indicate WT and HD distributions. **(C)** Transition probability matrices showing syllable-to-syllable transition structure in 12-month-old WT and HD mice, together with the WT-minus-HD subtraction matrix. Colored boxes indicate higher-order behavioral clusters. **(D)** Cluster-level transition networks showing genotype-associated changes in transition probability within slow rotation, excessive limb movement, segmented locomotion, and sedation clusters. Nodes represent syllables, arrows represent directed transitions, and line width indicates the absolute WT-HD transition probability difference. Data are presented as mean ± SEM where applicable.

We next examined pose-defined syllable distributions within representative HD-associated clusters with UMAP. WT and HD mice showed distinct density patterns within multiple significant behaviors, indicating that HD motor dysfunction reshaped the pose-dynamic organization of these behavioral states (**Figure 3B**). Syllable-to-syllable transition matrices further showed altered transition structures in 12-month-old HD mice. The WT-HD subtraction matrix revealed localized transition differences within and between higher-order behavioral clusters (**Figure 3C**).

Cluster-level transition networks were then generated to identify altered transition routes within specific behavioral clusters. Genotype-associated transition differences were observed within the 4 significantly changed behavior clusters (**Figure 3D**). These findings suggest that HD-associated behavioral impairments involve both altered occurrence of specific motor states and reorganization of the transition architecture connecting individual motor syllables.

### Behavioral flow analysis revealed progressive motor-state patterns across disease stages

To examine behavioral patterns across phenotypic progression, the common syllable-to-cluster framework was applied to WT and zQ175DN mice at 6, 9, and 12 months, from early phenotypic abnormalities, intermediate progression to pronounced motor impairment^11,30^. Top-view open-field recordings were analyzed to quantify cluster occurrence and transition structure, followed by age-specific construction of a mouse HD motor dysfunction score (**Figure 4A**).

**Figure 4.**
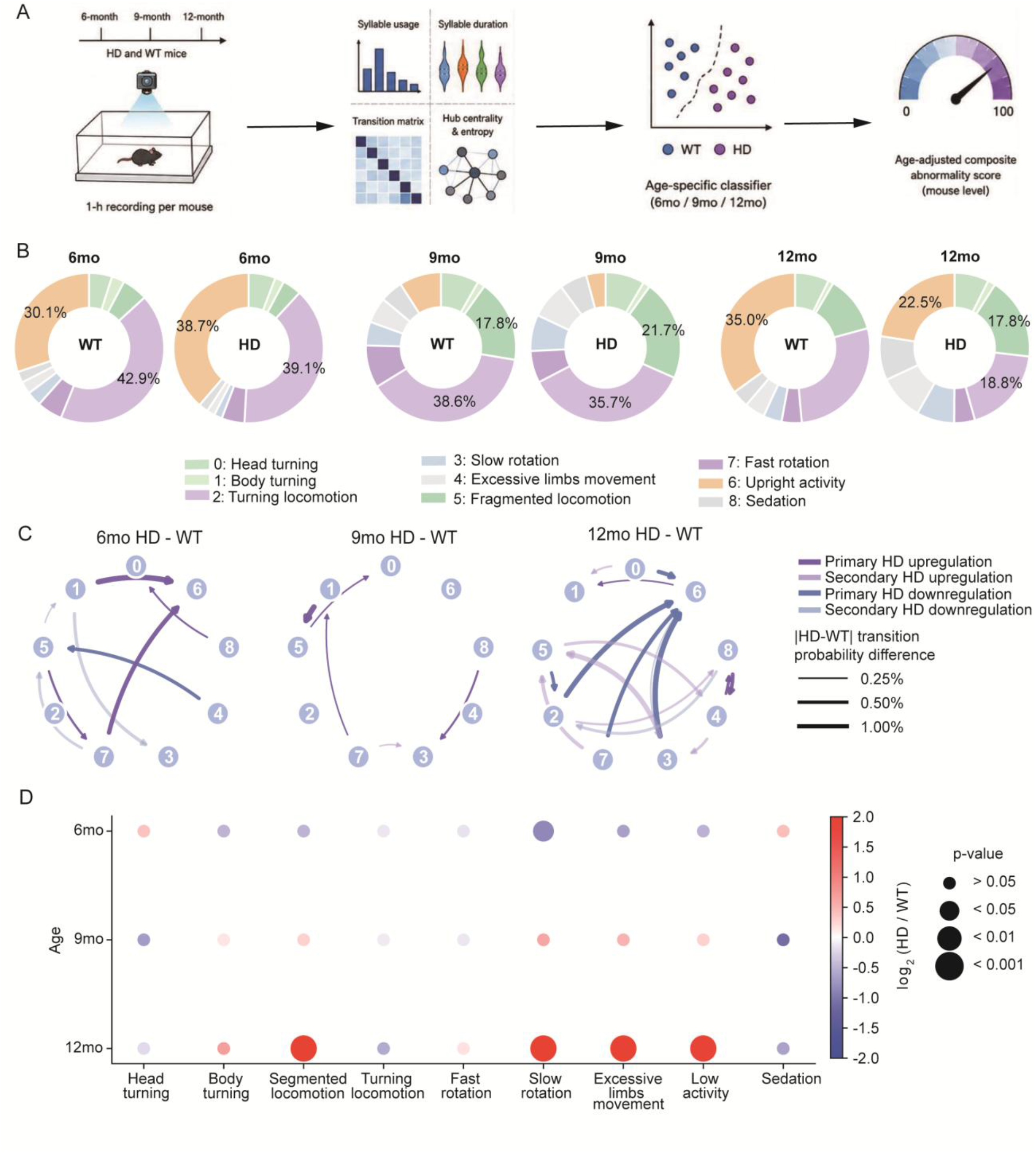
Behavioral flow analysis reveals age-dependent pattern changes of motor-state dynamics in HD mice. **(A)** Schematic overview of behavioral flow analysis. One-hour open-field recordings from WT and HD mice at 6, 9, and 12 months of age were analyzed to quantify age-dependent changes in behavior occurrence, integrated to define age-dependent motor-state pattern in HD mice. **(B)** Donut plots showing the occurrence composition of major behavioral clusters in WT and HD mice in 3 age groups. (**C)** Network plots showing genotype-associated differences in transition probability between behavioral clusters at 6, 9, and 12 months, presented as HD minus WT. Arrow color indicates the direction of HD-associated change, and line width indicates the magnitude of the transition probability difference. **(D)** Bubble plot summarizing age-dependent changes in cluster occurrence in HD mice relative to WT mice. Bubble color indicates log₂-transformed fold change, and bubble size indicates the corresponding p-value category.

The composition of higher-dimensional behavioral states differed across both age and genotype (**Figure 4B**). At 6 months, the overall behavioral repertoire remained broadly similar between WT and HD mice, although modest redistribution among upright activity, turning locomotion, and other motor states was evident. Greater genotype-dependent differences emerged at 9 months, including altered contributions of turning-related and locomotor states. By 12 months, HD mice showed a more pronounced redistribution of behavioral-state occurrence, with reduced contributions from upright activity and turning locomotion and increased contributions from segmented locomotion, slow rotation, excessive limb movement, and low-activity-related states.

Age-dependent motor patterns were also observed in the organization of transitions between behavioral clusters. HD-WT differential networks identified both accelerated and reduced transition routes at each age, as well as the affected connections across disease stages (**Figure 4C**). At 12 months, pronounced motor impairment affected a broader range of behavioral transitions.

Cluster-occurrence analysis further demonstrated that genotype effects were both stage-specific and behavior-dependent (**Figure 4D**). Developed HD was associated with prominent increases in segmented locomotion, slow rotation, excessive limb movement, and low-activity states. These findings indicate that HD progression involves multidimensional pattern changes of behavioral-state composition and transition architecture rather than a gross decline in spontaneous activity.

### Age-specific motor feature analysis identified distinct behavioral signatures in HD

To determine which behavioral feature domains are most distinguishable between HD and WT mice at different disease stages, we performed temporal-bin optimization and systematic feature-space screening. Linear discriminant analysis visualized age- and genotype-dependent structure across cluster bout frequency, cluster transitions, cluster occurrence, syllable bout frequency, combined features, syllable transitions, and syllable occurrence (**Figure S3A–G**). Classification performance varied with temporal resolution and disease stage. The F1 score is a classification performance metric that considers both the accuracy of positive predictions and the proportion of positive cases correctly identified. 15-minute bins provided the highest F1 score and accuracy in 9-month-old mice and maintained comparatively stable performance across ages; they were therefore selected for subsequent analyses (**Figure S3H,I**).

Receiver-operating characteristic analysis further showed that the most informative feature domain differed by age. Syllable mean duration produced the highest classification performance in 6 months, although with modest discrimination (AUC = 0.63). Syllable duration also performed best in 9 months, with improved discrimination (AUC = 0.81), whereas the combined feature space achieved the highest performance in 12 months (AUC = 0.85; Figure S3J). These results indicated that syllable-duration remodeling was the principal discriminative feature at early and intermediate stages, whereas a broader combination of behavioral dimensions was needed to characterize HD progression.

With the selected age-specific feature spaces, cross-validated confusion matrices showed limited WT–HD separation at 6 months but stronger classification at 9 and 12 months **(Figure 5A).** Accuracy increased from 0.57 (6 months) to 0.82 (9 months), and remained 0.77 at 12 months. Cross-validated predicted HD probabilities did not differ significantly between WT and HD mice at 6 months but were significantly higher in HD mice at 9 and 12 months **(Figure 5B).** Out-of-fold HD probabilities also increased across age in HD mice, with significant differences between 6 and 9 months and between 6 and 12 months, whereas no significant age-dependent changes were detected in WT mice (**Figure 5C**).

**Figure 5.**
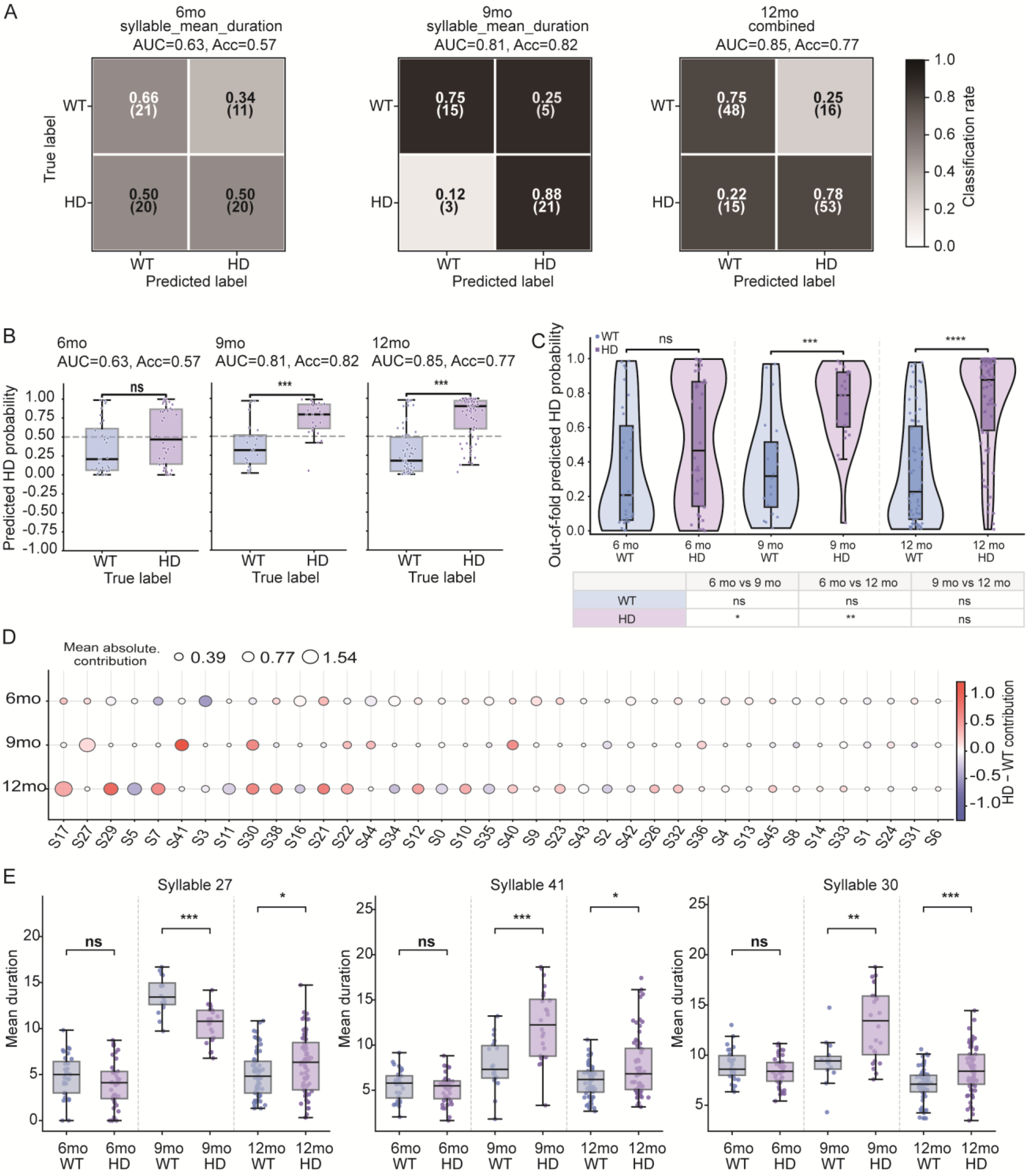
Stage-specific behavioral features classify HD mice across disease progression. At each disease stage, WT vs. HD mice was classified by behavioral features with ElasticNet-regularized logistic regression. GroupKFold cross-validation was grouped by mouse identity to prevent data leakage between temporal bins from the same mouse. **(A)** Confusion matrices for WT versus HD classification using the selected 15-min bin feature space at each age. **(B)** Predicted HD probability in WT and HD mice at 6, 9, and 12 months, showing stronger WT–HD separation at 9 and 12 months. **(C)** Age-dependent trajectory of predicted HD probability across 6-, 9-, and 12-month cohorts. **(D)** Duration-based feature contributions across disease stages. Dot color indicates the direction of HD–WT duration change, and dot size indicates mean absolute contribution to classification. **(E)** Mean duration of representative syllables showing age-dependent genotype effects. ns, not significant; *P < 0.05; **P < 0.01; ***P < 0.001.

Feature-contribution analysis identified distinct syllable-duration signatures across disease stages. Both positive and negative contributions were observed, indicating that HD classification reflected selective prolongation and shortening of individual syllables rather than a uniform directional change in bout duration (**Figure 5D**). Representative syllables exhibited this stage dependence: syllable 27 was shortened at 9 months and prolonged at 12 months, whereas syllables 41 and 30 were prolonged in HD mice at 9 and 12 months but were unchanged at 6 months **(Figure 5E)**. Together, these findings demonstrate that the behavioral features most informative for HD classification changed across disease progression, with selective syllable-duration remodeling emerging prominently at the intermediate stage and multidimensional behavioral disorganization characterizing developed HD.

### Transition-network remodeling as a dominant feature of developed HD

Because the combined feature space provided the strongest classification at 12 months, we examined the contribution of individual behavioral feature classes at this stage. Syllable-transition features were identified as the largest contributors to WT–HD separation, exceeding the contributions of occurrence, bout-frequency, duration, and cluster-level features (**Figure 6A**). This finding indicated that the developed HD phenotype was not just characterized by changes in individual motor states; it was also affected by reorganization of the transitions connecting them.

**Figure 6.**
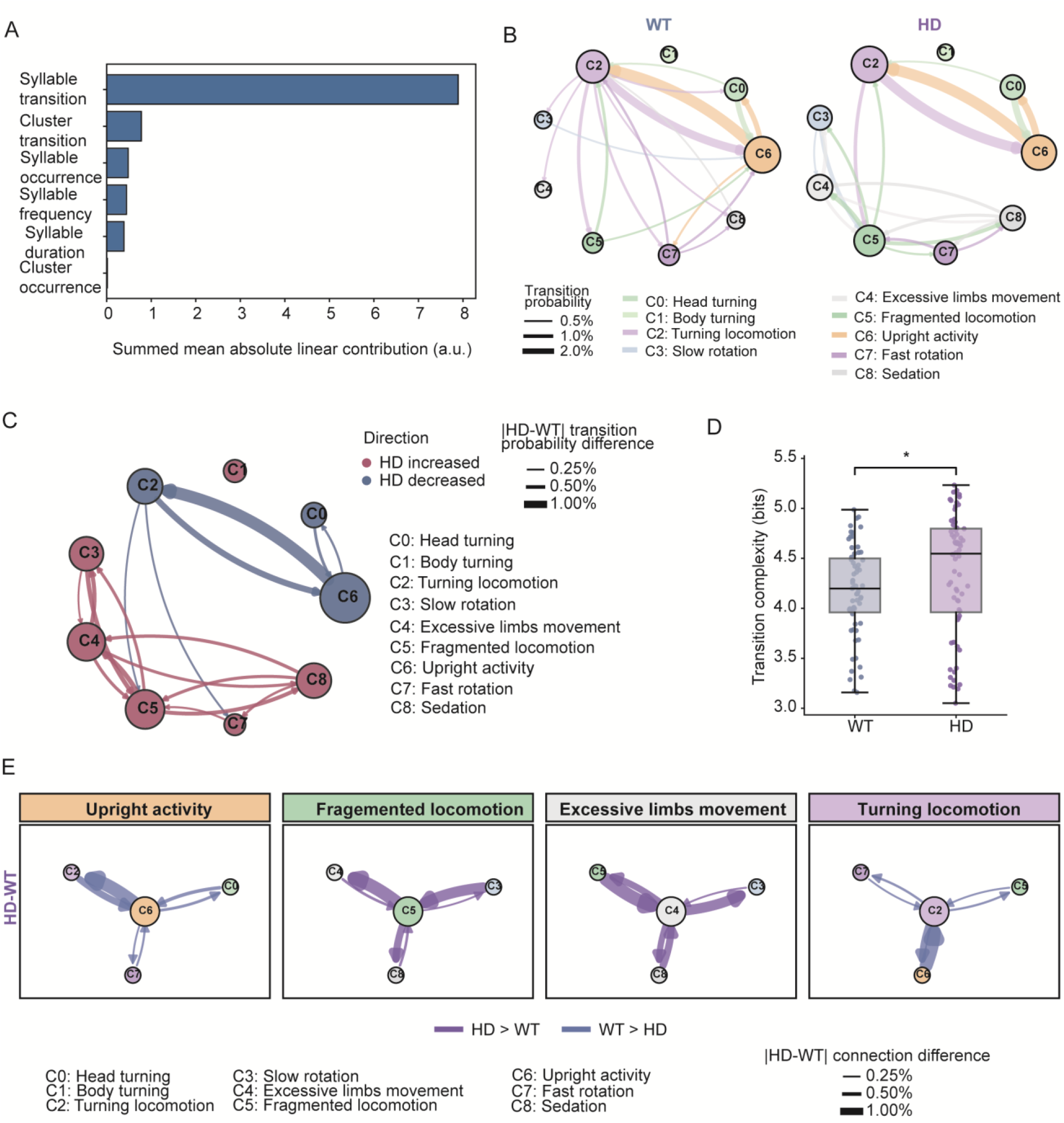
Late-stage HD is dominated by transition-network disorganization. **(A)** Feature-type contribution in the 12-month combined classifier, showing that syllable transition features were the strongest contributors to WT–HD separation. **(B)** Cluster hub networks in 12-month-old WT and HD mice. **(C)** HD–WT cluster transition hub network showing transitions increased or decreased in HD mice. **(D)** Transition complexity analysis showing increased behavioral transition complexity in 12-month-old HD mice. **(E)** Connection properties of representative behavioral hubs in WT and HD mice. *p < 0.05.

Genotype-dependent changes in the transition network were further elucidated by cluster-level hub analysis. Distinct hub configurations were observed in 12-month-old WT and HD mice, and both strengthened and weakened transition routes were identified in the HD-WT differential network (**Figure 6B,C**). These changes involved several locomotor and postural states and reflected redistribution of network centrality rather than a uniform increase or decrease in transition activity. Transition complexity was also increased in 12-month-old HD mice, indicating that behavioral transitions were distributed across a broader set of routes (**Figure 6D**). Connection-level analysis further confirmed bidirectional remodeling around representative behavioral hubs (Figure 6E).

Different transition-network abnormalities were observed at earlier disease stages. At 6 months, transition complexity was reduced in HD mice, whereas no significant genotype difference was detected at 9 months (**Figure S4A,B**). Despite these differences in global complexity, altered hub organization was already evident at both ages (**Figure S4C,D**). Local network analysis further showed that the direction and distribution of transition changes varied across disease stages (**Figure S4E,F**). Together, these findings indicate that transition-network remodeling emerged before pronounced motor impairment, but its organization and global complexity changed across HD progression, becoming a principal feature of the pronounced motor impairments.

### The HD motor dysfunction score provided a stage-sensitive endpoint

To integrate complementary behavioral abnormalities at the individual-mouse level, we constructed an HDMD score from syllable-duration remodeling, behavioral hub shift, and transition complexity (**Figure 7A**). We standardized each component against the age-matched WT distribution and used the absolute z-score to quantify the magnitude of deviation, regardless of direction. The HDMD score was then obtained as the mean of the three component scores, thereby providing an interpretable summary of abnormalities in motor-state persistence and transition-network organization.

**Figure 7.**
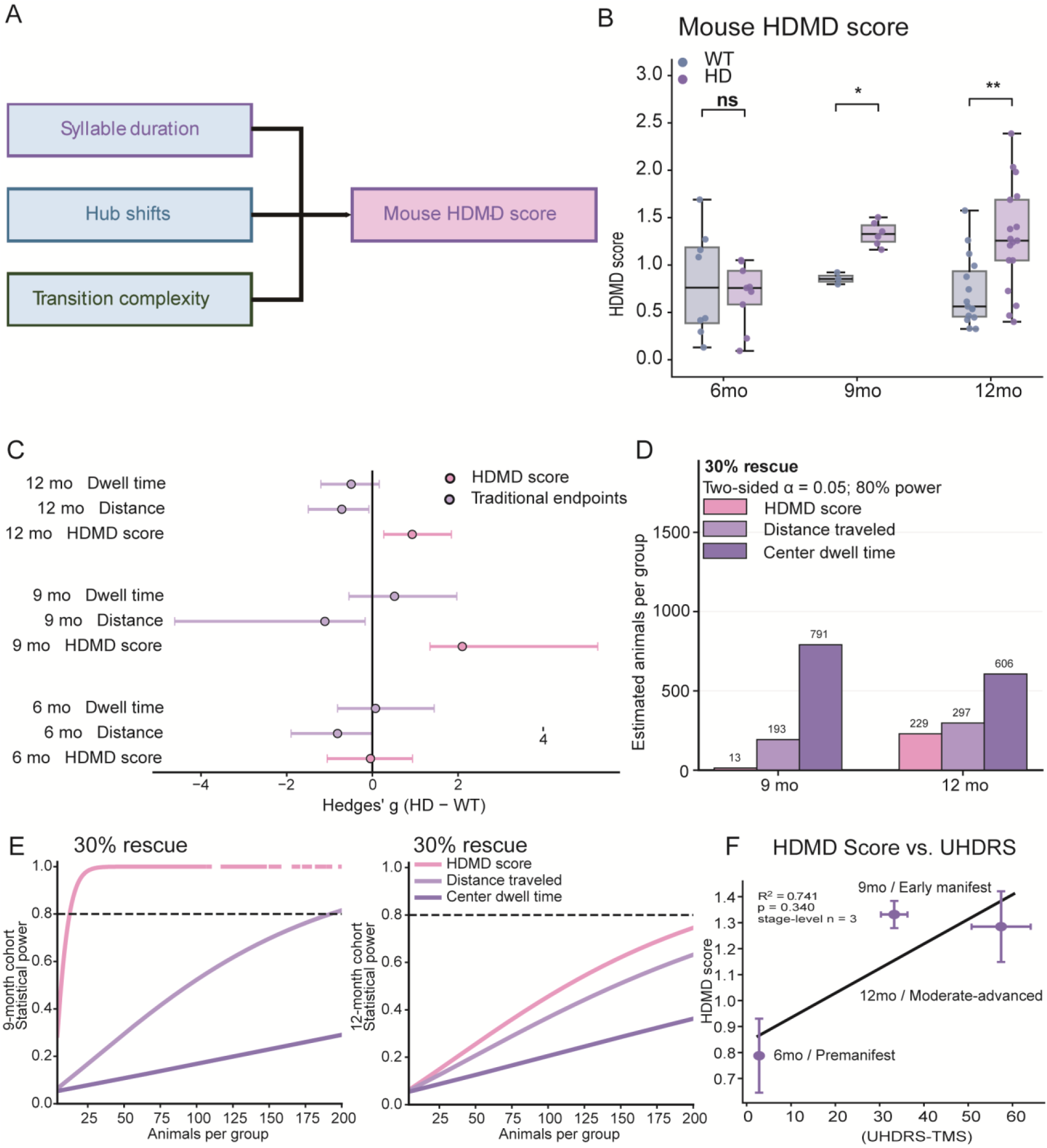
The HD motor dysfunction score captures stage-dependent motor abnormalities and improves statistical efficiency for detecting modeled therapeutic normalization. The HDMD score is the mean absolute deviation of three age-matched WT-standardized components: syllable duration, behavioral hub shift, and transition complexity. **(A)** HDMD score construction from age-matched WT-standardized absolute z-scores for syllable duration, hub shifts, and transition complexity. **(B)** HDMD scores in WT and HD mice at 6, 9, and 12 months. Box plots show the median and interquartile range, with whiskers indicating the full range; dots represent individual mice. WT and HD groups were compared at each age using two-sided Mann–Whitney tests. **(C)** Hedges’ g for HDMD score, distance travelled, and center dwell time; error bars indicate bootstrap 95% confidence intervals. **(D)** Estimated sample size per group required to detect a 30% normalization toward the age-matched WT mean at 80% power and two-sided α = 0.05. Estimates are shown for the HDMD score, distance travelled, and center dwell time at 9 and 12 months. **(E)** Statistical power curves for detecting a modeled 30% normalization at 9 months (left) and 12 months (right). The dashed horizontal line indicates 80% power. **(F)** Exploratory stage-level alignment between mouse HDMD score and human UHDRS total motor score. Points show stage means with error bars and a descriptive linear fit. P < 0.05, P < 0.01; ns, not significant.

Higher HDMD scores were detected in older HD mice with pronounced motor dysfunction. Also, no significant genotype difference was observed in 6-month-old HD mice (**Figure 7B**). Component-level analysis showed that syllable-duration remodeling was increased at 9 and 12 months (**Figure S6A**). Hub-shift abnormalities became significant at 12 months (**Figure S6B**). The age-matched WT-standardized transition complexity was not significantly increased, although a clear trend was observed at 12 months (**Figure S6C**). Thus, different behavioral features contributed to the HDMD score at different disease stages, rather than by uniform changes involving all components.

Next, we compared the performance of the HDMD score with traditional open-field test parameters. HD mice showed a gradual reduction in total distance traveled, whereas center dwell time was not significantly altered at all stages (**Figure S5A,B**). Representative occupancy maps illustrated differences in spatial exploration but did not provide a quantitative measure of fine-scale motor organization (**Figure S5C,D**). Hedges’ g estimates further showed that the relative sensitivity of each endpoint varied by disease stage (**Figure 7C**). A larger standardized genotype effect was obtained for the HDMD score than for traditional parameters at the progressing or pronounced motor impairment stages.

The implications for preclinical pharmacological study design were assessed by assessing partial normalization of the HD motor phenotype toward the age-matched WT mean. Under a 30% rescue assumption, 13 animals per group were estimated to be required using the HDMD score at 9 months, compared with 193 for distance traveled and 791 for center dwell time (**Figure 7D**).

At 12 months, the corresponding estimates were 229, 297, and 606 animals per group, respectively. Power analysis curves consistently showed that the HDMD score reached 80% power with markedly smaller cohorts at 9 months. HDMD scores also showed a modest advantage over distance at 12 months against traditional parameters (**Figure 7E**). These analyses indicate that the benefit of the composite endpoint was stage-progression dependent, where structured motor-flow abnormalities were detected despite comparatively limited changes in conventional measures.

To determine its translational value, we examined whether the HDMD score aligned with biological and clinical disease burden. Stage-level HDMD values increased in parallel with published human UHDRS total motor scores across corresponding disease stages (**Figure 7F**). Descriptive alignment was also observed with striatal mHTT aggregation, medium spiny neuron marker, and human CSF mHTT burden (**Figure S6D–G**). Despite limited disease stages in this study, these results were considered exploratory but still suggested that the behavioral analysis results indicate pathological and clinical disease severity.

Together, these findings establish the HDMD score as a stage-sensitive pharmacodynamic endpoint for HD preclinical research for the first time. Integrating motor-state features and transition-network abnormalities achieved higher HDMD sensitivity than traditional open-field parameters. This HDMD score provides a quantitative measure for endpoint selection, power optimization, and the design of future preclinical efficacy studies.

## Discussion

In this study, computer vision-aided behavioral flow analysis revealed stage-specific changes in spontaneous locomotion organization in zQ175DN mice. Conventional open-field measures captured reduced locomotion at symptomatic stages but provided limited information about how individual motor states were expressed over disease progression. In contrast, bidirectional changes were detected in behavioral syllables, indicating that HD motor dysfunction was not caused by uniform behavioral suppression, which may reflect both disease progression and compensatory adjustment in different disease stages. Alterations were also found in syllable duration, higher-order behavioral states, and transition networks. These findings extend earlier studies showing that subtle neural and behavioral abnormalities can emerge before robust changes are detected by conventional motor tests^11,31^.

The behavioral features that best distinguished HD mice from WT differed across disease stages. Syllable-duration remodeling provided the strongest discrimination at earlier and intermediate stages, whereas transition features became more informative in more advanced stages. Hub organization and transition complexity were also altered across stages, but the direction of these changes was not uniform. Transition complexity was reduced at the early stage and increased in developed HD. This pattern suggests that HD progression is associated with changing forms of motor disorganization rather than a simple linear loss of behavioral complexity. Spontaneous behavior is organized across multiple temporal levels, with motif duration and transition structure providing distinct information^20,21,32^. Region-specific neurotransmitter dynamics may also encode behavior across different timescales, supporting the biological relevance of syllable-duration changes^33^.

The higher-order behavioral clusters improved the interpretation of individual syllables. Developed HD was associated with increased segmented locomotion, slow rotation, excessive limb movement, and low-activity-related states. These changes were supported by several syllables within each cluster and were therefore not driven by isolated motifs. Hierarchical behavioral frameworks offer a useful bridge between short motor motifs and broader behavioral sequences, but post hoc annotation remains an important source of uncertainty^29^.

The stage-specific behavioral signatures are consistent with progressive corticostriatal dysfunction in HD. Striatal circuits contribute to the selection, maintenance, and switching of motor actions^22,34,35^. Altered syllable duration may therefore reflect impaired maintenance or termination of individual motor programs. Hub remodeling and dispersed transition routes may instead reflect impaired organization of motor sequences. This interpretation is also consistent with stage-dependent molecular changes in corticostriatal cells and the selective vulnerability of medium spiny neurons^3,4^. Corticostriatal dysfunction in HD also involves altered astrocytic engagement of striatal synapses, which may further disturb the control of motor-state transitions^5^. These links remain indirect because neural activity was not measured in the present study. Simultaneous behavioral and circuit recording will be needed to determine which neural changes drive each behavioral component.

The HDMD score was developed to summarize these complementary abnormalities at the individual-mouse level. It combined age-matched WT-standardized deviations in syllable-duration remodeling, behavioral hub shift, and transition complexity. Unlike a black-box classifier, each component represents a defined aspect of behavioral organization^20^. The score also captures bidirectional abnormalities, because a feature can indicate motor dysfunction when it is either higher or lower than in same-stage WT mice. HDMD distinguished WT and HD mice at intermediate and advanced stages. It should therefore be viewed as a stage-sensitive measure rather than a universal endpoint for all phases of HD.

Another contribution of this study is an interpretable framework for detecting therapeutic normalization of motor organization. A candidate treatment may improve the persistence or sequencing of motor states without fully restoring total distance traveled. Such effects may be missed when efficacy is judged only from gross locomotor activity. This is relevant to HD drug development, where treatments with different mechanisms may produce distinct functional response profiles^36^. In contrast, increased locomotion alone does not necessarily indicate therapeutic benefit and may instead reflect nonspecific psychostimulant or other motor effects^37^. By separating overall activity from behavioral structure, the HDMD score may help distinguish organized recovery from nonspecific motor activation.

Unlike earlier high-resolution HD phenotyping studies designed primarily for genotype classification or early abnormality detection, the present framework was developed around preclinical pharmacological use^25^. The stage-specific features were integrated into an interpretable HDMD score, and its ability to detect modeled partial normalization was compared with conventional open-field endpoints. Moreover, using standard top-view recordings reduces hardware requirements and may allow existing treatment cohorts to be reanalyzed.

This distinction may be relevant to therapies with different mechanisms. Huntingtin-lowering interventions are expected to act upstream and may gradually normalize several behavioral dimensions as disease progression is slowed. Huntingtin silencing has already been shown to delay behavioral and biomarker progression in HD models^7^. In contrast, treatments that modify corticostriatal transmission may produce more selective changes in syllable persistence or transition-network structure. Manipulation of VGLUT3-dependent signaling, for example, can improve motor deficits and neuronal survival in zQ175 mice^38^. Behavioral-flow analysis could help determine whether such interventions restore specific motor-state relationships even when conventional locomotor recovery is incomplete. These applications remain hypothetical because the present study included no treatment cohort.

The component structure of the score may also help interpret differences among candidate compounds. A treatment that mainly reduces the syllable duration-remodeling component may act differently from one that restores hub organization^24^. This information could support compound prioritization when two treatments produce similar changes in total activity. It may also help identify the disease stage at which a mechanism has its strongest functional effect.

However, the total HDMD score should not be reported alone in treatment studies. The signed values of its individual components should also be examined. This is necessary to confirm that a lower composite score reflects movement toward the WT state rather than unrelated changes in the opposite direction.

The power analysis further supports a stage-specific approach to preclinical study design. At the intermediate stage, the HDMD score was estimated to detect 30% normalization with substantially fewer animals than distance traveled or center dwell time. Its advantage was smaller in developed HD. The best endpoint may depend on treatment timing, with syllable-duration or composite measures favoring earlier intervention and transition-network features favoring developed HD. These estimates were derived from untreated WT and HD distributions. They do not represent prospective pharmacological validation and may change if treatment affects variability, exposure, attrition, or the shape of the response.

Prospective validation is recommended before using the HDMD score as a primary efficacy endpoint. Independent treatment cohorts should include a benchmark intervention with established target engagement and behavioral benefit. Dose-response and exposure-response relationships should also be examined. A valid pharmacodynamic endpoint should change in a biologically plausible direction and should track target engagement or downstream pathology when appropriate^39^. Pharmacological challenge studies will also be important. Sedative, stimulant, anxiolytic, and stereotypy-inducing treatments may alter syllable duration or transition complexity without correcting HD pathology. Including compounds with known motor liabilities would help define these nonspecific signatures and test whether HDMD improvement reflects genuine normalization^40^.

A practical advantage of the present framework is that it requires only standard top-view open-field recordings. More complex multi-camera and depth-sensing systems can provide better information about vertical posture and fine limb movement, but such systems are not routinely used in many pharmacological studies. Keypoint-MoSeq and pretrained pose models can be applied to conventional laboratory videos, creating an opportunity to extract additional information from existing recordings^20,26^. Historical treatment cohorts could therefore be reanalyzed without repeating the original animal experiment. This may support comparison of compounds, doses, and treatment windows while reducing cost and additional animal use.

The use of archived recordings will nevertheless require careful qualification. Pose estimation and syllable segmentation can be affected by frame rate, image resolution, camera position, arena geometry, lighting, and body-point visibility. Behavioral-video models may capture both disease-related signals and unrelated variation arising from subjects or recording conditions^41^. A model trained on one recording system may not generate directly comparable scores in another setting. The current workflow should therefore be tested across laboratories and recording conditions before establishing a shared reference range. Locked feature definitions, predefined quality-control thresholds, and double-blinded analysis will also be needed to limit analytical flexibility ^26,42^. The use of same-stage control may further require concurrent control groups or a well-qualified normative dataset.

The exploratory pathological and clinical comparisons provide initial support for the biological relevance of the HDMD score. Stage-level increases in HDMD were aligned with greater mHTT burden, reduced medium spiny neuron survival, and higher human motor severity. UHDRS motor measures have previously been associated with progression in imaging biomarkers, while CSF mHTT reflects molecular disease burden in patients^43,44^. The zQ175DN model also shows progressive metabolic impairment and biomarker changes across disease stages^30^. However, only three stage-level observations were available for these comparisons. They do not establish individual-level correlation, prediction, or clinical validity. Future studies should relate mouse-level HDMD scores to pathology measured in the same animals and determine whether similar behavioral-flow features can be captured in patients using digital motor assessments.

There are several limitations in the current study. The behavioral vocabulary was derived from a late-stage discovery cohort and then applied to all stages. This provided a shared feature space but may have underrepresented motifs that occur mainly during earlier disease. The cohorts were cross-sectional, so the apparent shift from duration remodeling to network disorganization cannot be interpreted as a within-animal sequence. The intermediate-stage sample was also relatively small, which may increase uncertainty in classification, effect-size, and power estimates. HD behavioral phenotypes are affected by sex, body weight, and testing conditions, and these factors should be examined in larger independent cohorts^11,12^. Finally, the framework was limited to spontaneous behavior in an open field. It does not replace tests of coordination, balance, cognition, or other HD-affected functions.

In conclusion, HD progression was characterized by structured changes in motor-state persistence and transition-network organization rather than a uniform decline in movement. Distinct behavioral features emerged across disease stages and were integrated into an interpretable HDMD score. The score showed sensitivity to partial rescue at the intermediate stage and may improve endpoint selection in preclinical studies. Its compatibility with standard top-view videos also enables existing treatment datasets to be revisited at higher behavioral flow. Together, these features position the HDMD framework as a practical and scalable strategy for pharmacodynamic assessment in HD drug development.

## Methods

### Animals

Heterozygous zQ175DN knock-in mice were originally obtained from The Jackson Laboratory (Bar Harbor, ME, USA; Stock No. 029928) and maintained as an in-house breeding colony at Johns Hopkins University. Mice were bred according to the established breeding protocol to generate heterozygous zQ175DN mice and wild-type (WT) littermate controls. WT and heterozygous zQ175DN mice were studied at 6, 9, and 12 months of age. Male and female mice were included at an approximately 1:1 ratio. Body weights at the time of study ranged from approximately 20 to 35 g.

Group sizes were as follows: 6 months, WT (n = 8) and HD (n = 10); 9 months, WT (n = 5) and HD (n = 6); and 12 months, WT (n = 16) and HD (n = 17). Animals were housed under standard conditions maintained by the Johns Hopkins University Research Animal Resources (RAR), with ad libitum access to food and water.

All animal procedures were approved by the Johns Hopkins University Animal Care and Use Committee (ACUC; protocol no. MO24M120) and were performed in accordance with institutional guidelines for the care and use of laboratory animals.

### Open-field behavioral recording

Mice were placed individually into an open-field arena and recorded for 1 h. Videos were acquired from a top-view camera positioned above the arena. The arena size was 40cm×40cm, and videos were recorded at 30 frames or 8 frames per second under standardized lighting conditions. The 1-h recording was applied for both conventional behavioral analysis and computational behavioral segmentation.

Conventional open-field parameters are tracked with a self-trained YOLO model, including total distance traveled, average speed, dwell time in the arena center, and time per visit in the middle zone. Distance and speed were applied as measures of spontaneous locomotor output, whereas center-zone measures were applied to assess spatial exploration. Spatial occupancy maps were generated by accumulating animal positions across the recording session and visualizing the relative time spent in each arena location.

### Pose tracking and estimation

Mouse pose was estimated from top-view open-field videos using DeepLabCut Model Zoo. We applied the SuperAnimal-TopViewMouse pretrained model family, which is designed for laboratory mouse videos acquired from a top-view perspective. Pose inference was performed through the DeepLabCut graphical interface using the superanimal_topviewmouse supermodel with the PyTorch engine. The pose estimator was set to resnet_50, and the detector was set to fasterrcnn_mobilenet_v3_large_fpn. Because each open-field recording contained a single mouse, the maximum number of individuals was set to 1.

For inference, the pose confidence threshold was set to 0.40, and the detector confidence threshold was set to 0.10. The pose model batch size and detector batch size were both set to 1. Self-supervised video adaptation was enabled to improve tracking stability on the open-field recordings, with a pseudo-label confidence threshold of 0.10, 4 pose adaptation epochs, and 4 detector adaptation epochs. Labeled videos were generated and visually inspected for quality control before downstream Keypoint-MoSeq behavioral segmentation.

### Keypoint-MoSeq behavioral segmentation

Keypoint-MoSeq was applied to segment open-field behavior into recurrent behavioral syllables^20^. Because 12-month-old HD mice showed the most pronounced motor phenotype, this age was applied as the discovery stage to define the behavioral syllable vocabulary. For model training, we selected a balanced subset of four 12-month recordings, consisting of two WT and two HD mice. DeepLabCut .h5 key point files from these recordings were applied as input to Keypoint-MoSeq.

The model was initialized using a PCA representation of the formatted pose data. The location noise parameter was estimated from the pose trajectories using a filtering scale based on the recording frame rate. Model fitting was performed in two stages. First, an autoregressive-only model was trained for 50 iterations with jitter set to 0.01. The resulting checkpoint was then applied to initialize full model fitting. Before full model training, (*κ*)was set to 1 × 10^2^. The full Keypoint-MoSeq model was then trained for an additional 100 iterations with jitter set to 0.05. After model fitting, syllables were reindexed to generate a consistent syllable order for downstream analyses.

The trained model identified 94 behavioral syllables, each representing a short recurrent movement motif defined from pose dynamics. This 12-month-derived syllable flow architecture was applied as a common behavioral vocabulary for subsequent analyses and was applied across 6-, 9-, and 12-month WT and HD cohorts. Syllable usage was calculated as the fraction of frames assigned to each syllable, syllable bout frequency as the number of syllable bouts normalized by recording duration, syllable mean duration as the average duration of bouts assigned to each syllable, and syllable transition probability as the probability of transitions between consecutive syllable labels.

### Cluster tree build

To identify higher-order behavioral states, Keypoint-MoSeq syllables were grouped according to their outgoing transition probability profiles. Hierarchical clustering was performed to group syllables according to their temporal organization within the behavioral sequence. Ward linkage with Euclidean distance was applied to cluster syllables with similar transition profiles. The resulting dendrogram was cut to generate nine target communities. Community identities were then reordered according to their left-to-right order in the dendrogram to produce consistent cluster labels for downstream visualization and analysis.

To avoid treating very small or rarely expressed branches as stable behavioral states, communities were filtered based on branch size and total bout count. Communities containing fewer than two syllables or fewer than 80 total bouts were labeled as small/outlier communities and assigned to the outlier category. The remaining communities were retained as stable behavioral clusters.

This transition-profile-based syllable-to-cluster mapping allowed individual syllables to be summarized into higher-order behavioral state classes. Cluster usage was then calculated as the summed usage of all syllables assigned to each cluster. Cluster transition features were calculated by converting syllable labels into cluster labels and quantifying transitions between consecutive cluster states. The resulting cluster-level representation was applied to analyze behavioral state composition, cluster-level transition structure, hub centrality, and transition complexity.

### Temporal-bin optimization and age-specific feature-space screening

To identify the temporal resolution and behavioral feature domain that best separated WT and HD mice at each disease stage, we performed systematic bin-level classification. For each temporal window size, age, and feature block, WT-versus-HD classifiers were trained using ElasticNet-regularized logistic regression. The classification pipeline included variance filtering, z-score standardization, and logistic regression with ElasticNet penalty. The logistic regression model applied the saga solver, l1_ratio = 0.5, C = 1.0, balanced class weights, and a maximum of 8,000 iterations.

To prevent data leakage across temporal bins from the same mouse, model evaluation applied GroupKFold cross-validation with mouse identity as the grouping variable. The number of folds was set to the smaller of five or the number of mice available for that age group. For each held-out fold, predicted HD probabilities were generated using cross-validated prediction.

Model performance was evaluated using ROC AUC, accuracy, balanced accuracy, HD precision, HD recall, HD F1 score, and confusion matrices. Because classifier score orientation can vary, raw AUC values below 0.5 were reoriented by reversing the predicted probability, yielding an oriented AUC. The best feature block for each age and temporal window was selected by maximizing oriented AUC, with balanced accuracy and HD F1 score applied as secondary ranking criteria. Fifteen-minute bins provided robust performance across ages and were selected for downstream bin-level classification and interpretation analyses.

### Model interpretation and feature contribution analysis

To identify behavioral features contributing to genotype classification, feature contribution analyses were performed for the best-performing age-specific classifiers. For linear models, feature contributions were derived from model coefficients and feature values after preprocessing. Features were ranked by mean absolute contribution to classification. The direction of contribution was applied to determine whether each feature contributed toward the HD or WT class. For 6- and 9-month classifiers, syllable mean-duration features were examined in detail. For the 12-month classifier, feature-class contributions were summarized to evaluate the relative contribution of usage, duration, frequency, transition, and cluster-level features.

### HDMD score

To summarize HD-associated behavioral disorganization at the mouse level, we constructed a composite chorea-like behavioral disorganization score using three interpretable behavioral components: a syllable-duration remodeling score, a behavioral hub shift score, and transition complexity. These components were selected to capture the major behavioral dimensions identified in the preceding analyses. Specifically, syllable-duration features were included because feature-space screening identified syllable mean duration as the most informative feature block for distinguishing WT and HD mice at 6 and 9 months. Hub shift and transition complexity were included to represent the transition-network abnormalities that became prominent at 12 months.

All score components were first calculated at the level of one mouse by one 15-min bin. The 15-min bin-level feature matrix was applied because temporal-bin optimization identified 15-min bins as a robust temporal resolution for genotype classification while preserving within-session behavioral structure.

### Duration remodeling score

The duration remodeling score was designed to quantify the syllable-duration signature identified from the early-to-mid-stage HD classifiers. This component was calculated from selected syllable mean-duration features rather than cluster-level duration features, because the most informative duration-based feature space for 6- and 9-month classification was the syllable mean-duration feature block.

Syllables were divided into HD-longer and HD-shorter sets based on the duration effect trajectory analysis. The HD-longer set included syllables 41, 30, 44, 33, 2, and 40. The HD-shorter set included syllables 22, 27, 24, and 36. For each selected syllable, the corresponding syllable mean-duration feature was z-scored across all analyzed 15-min bins. For each bin, the duration remodeling score was calculated as:

Higher values indicate stronger expression of the HD-associated syllable-duration remodeling pattern.

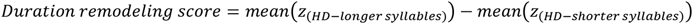

### Behavioral hub shift score

The behavioral hub shift score was designed to quantify whether the transition-network organization shifted toward HD-associated behavioral clusters and away from WT-associated clusters. For each 15-min bin, cluster-to-cluster transition features were applied to construct a directed transition matrix. Rows represented source clusters and columns represented target clusters. The transition matrix was row-normalized so that outgoing transition probabilities from each source cluster represented the relative transition distribution from that cluster.

The row-normalized transition matrix was then converted into a directed weighted graph, in which nodes represented behavioral clusters and directed edges represented cluster-to-cluster transition probabilities. PageRank centrality was calculated for each cluster to estimate its relative importance within the behavioral transition network.

HD-associated clusters were defined as C3, C4, C5, and C8. WT-associated clusters were defined as Rear/C9 and C2, depending on the cluster labels available in the feature matrix. For each 15-min bin, the hub shift score was calculated as:

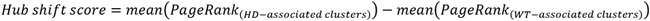

Higher values indicate a network-level shift toward HD-associated behavioral hubs.

### Transition complexity

Transition complexity was applied to quantify the complexity of behavioral state transitions. For each 15-min bin, all nonzero cluster-to-cluster transition probabilities from the transition matrix were collected and normalized to sum to one. Shannon entropy was then calculated as:

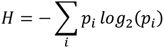

where *p_i_* represents the normalized probability of transition (i). Higher transition complexity (entropy) indicates greater transition complexity and a less concentrated transition structure.

### Bin-level composite HDMD score

After calculating the duration remodeling score, hub shift score, and transition complexity for each 15-min bin, each component was z-scored across all analyzed bins. The bin-level chorea-like behavioral dysfunction score was calculated as:

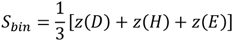

where (*S_bin_*)is the bin-level HDMD score, (D) is the duration remodeling score, (H) is the hub shift score, and (E) is transition complexity.

A transition-network-only score was also calculated as:

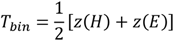

where (*T_bin_*)is the bin-level transition-network score. These bin-level scores were applied to summarize behavioral dysfunction within individual temporal segments of the open-field recording.

### Mouse-level score aggregation

To generate mouse-level scores for pathology correlation and group-level comparison, bin-level component scores and composite scores were averaged across all 15-min bins from the same mouse. This produced one mouse-level value for each variable, including the duration remodeling score, hub shift score, transition complexity, chorea-like score, and transition-network score.

### Age-adjusted HD motor dysfunction score

Since open-field behavior and behavioral state organization may change with normal aging, we further normalized the mouse-level behavioral components relative to age-matched WT baselines. This generated an age-adjusted chorea-like behavioral abnormality score designed to quantify disease-associated deviation from normal same-age behavioral organization.

For each age group and each mouse-level component, WT mice were applied to calculate the age-matched WT baseline mean and standard deviation. Each mouse was then assigned an age-matched WT-normalized z score for each component:

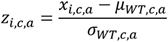

where (*x_i_*_,*c*,*a*_)is the value of component (c) for mouse (i) at age (a), and (*μ_WT_*_,*c*,*a*_) and (*σ_WT_*_,*c*,*a*_) are the mean and standard deviation of the same component in age-matched WT mice.

The three components applied for age adjustment were the mouse-level duration remodeling score, hub shift score, and transition complexity. To quantify the magnitude of behavioral abnormality regardless of whether a component increased or decreased, the absolute value of each age-matched WT-normalized z score was calculated. The final age-adjusted chorea-like behavioral abnormality score was defined as:

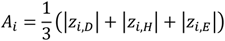

where (*A_i_*) is the age-adjusted chorea-like behavioral abnormality score for mouse (i), (*z_i_*_,*D*_)is the age-matched WT-normalized z score for the duration remodeling component, (*z_i_*_,*H*_) is the age-matched WT-normalized z score for the hub shift component, and (*z_i_*_,*E*_) is the age-matched WT-normalized z score for the transition complexity component.

This score measures how strongly each mouse deviates from the behavioral organization of same-age WT controls across the major behavioral dimensions identified by the classification and network analyses. Higher values indicate greater behavioral abnormality relative to the age-matched WT baseline.

For statistical comparison, WT and HD mice were compared within each age group. Data-exclusion criteria were defined before statistical testing and applied consistently across groups. Where indicated, outliers were identified using the 1.5×interquartile range (IQR) rule. Group comparisons were performed using two-sided Mann–Whitney U tests.

### Immunofluorescence staining and imagnig

Mice were deeply anesthetized with isoflurane (Fluriso, Cat. No. V1-502017) and transcardially perfused with phosphate-buffered saline (PBS, pH 7.4; Gibco, Cat. No. 10010031), followed by 4% paraformaldehyde (PFA; Sigma-Aldrich, Cat. No. P6148-500G). Brains were removed and post-fixed in 4% PFA overnight at 4 °C, followed by cryoprotection in 30% sucrose (Sigma-Aldrich, Cat. No. S0389-500G) in PBS. Tissues were embedded in OCT Tissue Freezing Medium (Leica Biosystems, Cat. No. 14020108926) and coronally sectioned at 30 μm thickness.

Free-floating brain sections containing the striatum were washed in PBS and permeabilized with 0.3% Triton X-100 (Sigma-Aldrich, Cat. No. T9284-100ML). Sections were blocked for 1 h at room temperature in PBS containing 5% normal goat serum (Fisher Scientific, Cat. No. NC1899949) and then incubated overnight at 4 °C with the appropriate primary antibodies. Primary antibodies included rabbit monoclonal anti-mHtt P90, clone 1B12 (CHDI Foundation, CHDI-90004290), rat monoclonal anti-NeuN (Abcam, Cat. No. ab279297), and rabbit monoclonal anti-DARPP-32 (Abcam, Cat. No. ab40801).

After washing with PBS, sections were incubated for 1 h at room temperature with the appropriate fluorophore-conjugated secondary antibodies protected from light. Secondary antibodies included goat anti-rabbit IgG (H+L) Alexa Fluor 488 (Invitrogen, Cat. No. A-11008), goat anti-rabbit IgG (H+L) Alexa Fluor 555 (Invitrogen, Cat. No. A-21428), and goat anti-rat IgG (H+L) Alexa Fluor 647 (Invitrogen, Cat. No. A-21247). Nuclei were counterstained with Hoechst 33342 (Sigma-Aldrich, Cat. No. 14533-100MG).

Fluorescence images were acquired using a Zeiss LSM 800 confocal microscope. Images used for quantitative comparisons were acquired using identical imaging settings within each experiment. Background correction was performed using ZEN, and quantitative image analysis was performed using QuPath. P90-mHTT fluorescence and neuronal-marker signals were quantified within predefined striatal regions of interest using identical analysis parameters across experimental groups. The mean value obtained from each animal was applied as one biological replicate for statistical analysis.

### Human HD-CSF dataset and cross-species stage-level comparison

Publicly available human clinical and biomarker data were obtained from the UCL Research Data Repository HD-CSF study mHTT and NfL dataset at DOI: 10.5522/04/12709391. This dataset was applied only for secondary stage-level comparison, and no new human data were collected in the present study. The corresponding HD-CSF study was cited as the original clinical biomarker study 44.

To assess cross-species disease relevance, mouse age groups were aligned conceptually with ordered human HD stages according to disease severity. Mean age-adjusted HD-like chorea scores from 6-, 9-, and 12-month zQ175DN mice were compared with stage-level human UHDRS total motor scores and CSF mHTT burden. Associations were assessed using linear regression and interpreted as exploratory stage-level comparisons rather than direct clinical validation.

### Quantification and statistical analysis

Data are shown as mean ± SEM unless otherwise stated. Each dot represents one mouse, one temporal bin, or one behavioral sample as indicated in the corresponding figure legend. Statistical analyses were performed using Python, GraphPad Prism, and SPSS.

For two-group comparisons, WT and HD mice were compared using Student’s t test, Welch’s t test, or Mann–Whitney U test, depending on data distribution and variance assumptions. For syllable-, cluster-, and transition-level analyses, outliers were removed within each feature × genotype group using the 1.5×IQR rule before statistical testing when indicated. Multiple comparisons were corrected using the Benjamini–Hochberg false discovery rate procedure when applicable. Statistical significance was defined as p < 0.05. Significance levels are indicated as ns, not significant; *p < 0.05; **p < 0.01; ***p < 0.001.

For classification analyses, performance was evaluated using GroupKFold cross-validation grouped by mouse identity. Classifier performance metrics included oriented ROC AUC, accuracy, balanced accuracy, precision, recall, F1 score, and confusion matrices. The best feature space was selected by oriented AUC, with balanced accuracy and HD F1 score applied as secondary criteria.

### Software and algorithms

Behavioral tracking was performed using self-trained YOLO and DeepLabCut. Behavioral segmentation was performed using Keypoint-MoSeq. Data processing, feature extraction, classifier training, transition-network analysis, and figure generation were performed in Python using NumPy, pandas, scikit-learn, SciPy, statsmodels, matplotlib, and network analysis packages. Statistical plotting was performed using Python and GraphPad Prism. Image analysis was performed using QuPath and ZEN.

## Acknowledgements

This project is supported by NIH R01 NS127344 and NIH R01 NS124084.

---

**Figure S1.**
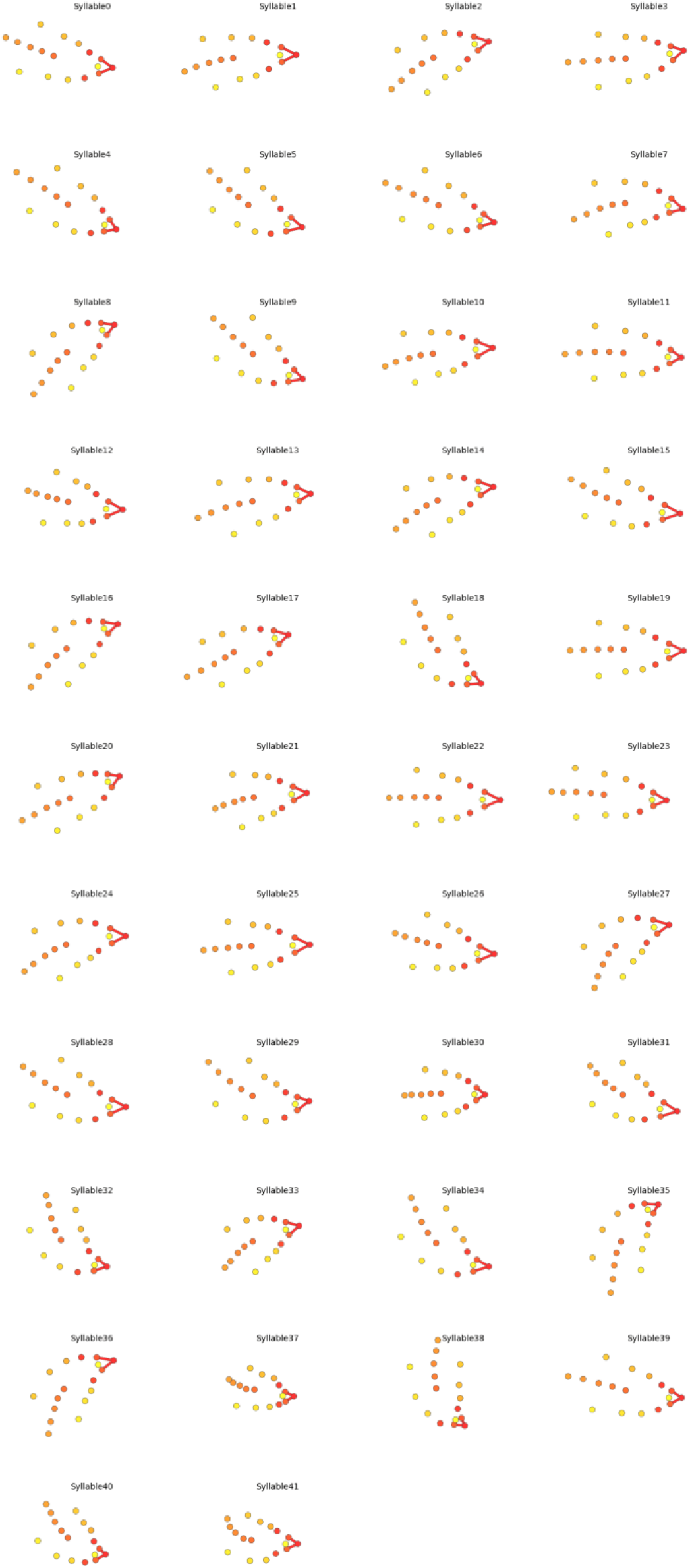
Representative pose trajectories of the 42 most frequently expressed syllables among the 45 syllables retained for downstream analysis.

**Figure S2.**
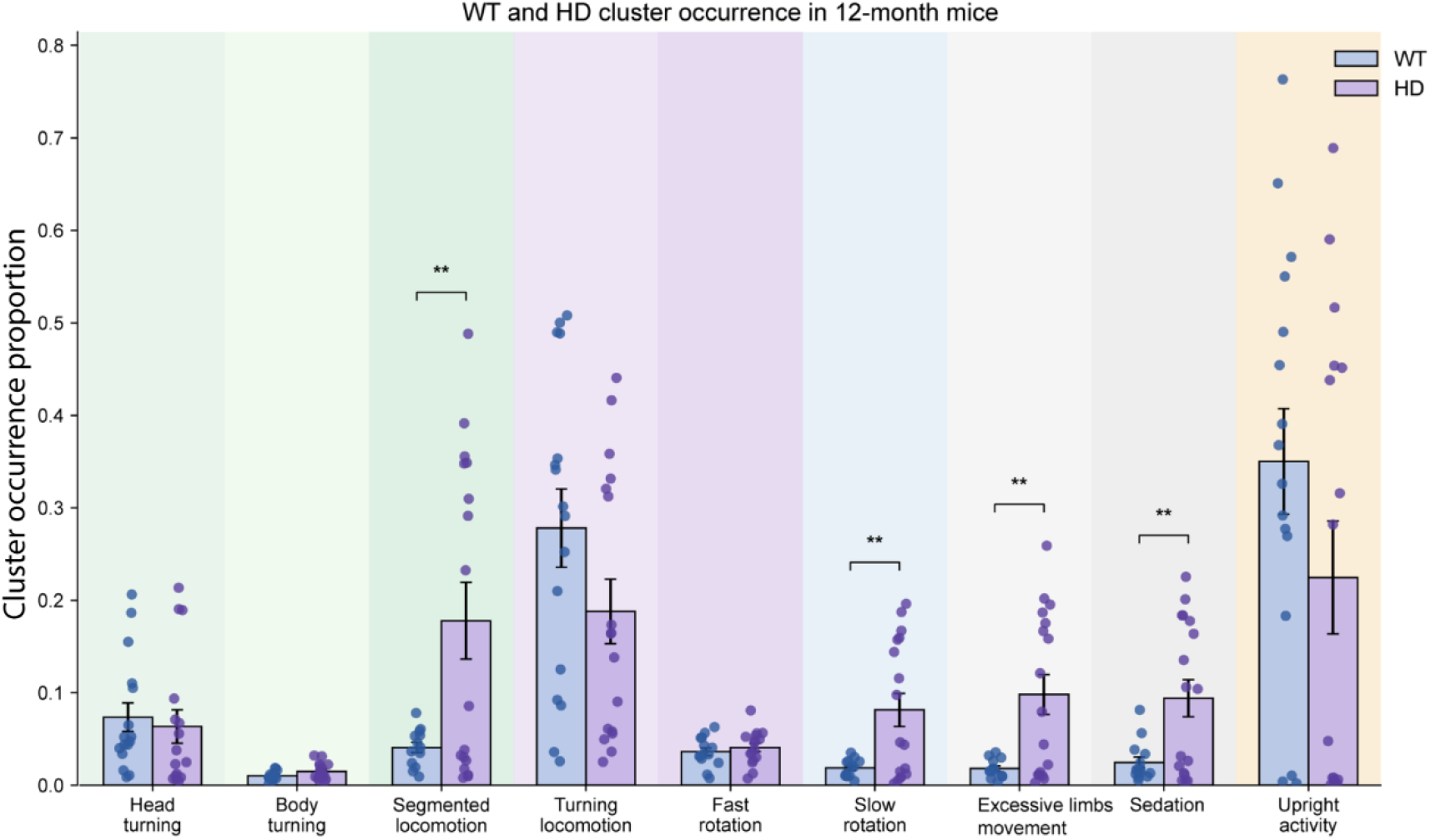
Cluster occurrence in 12-month-old WT and HD mice. Occurrence of major behavioral clusters in 12-month-old WT and zQ175DN HD mice. Each point represents one mouse, and bars show mean ± SEM. **P < 0.01.

**Figure S3.**
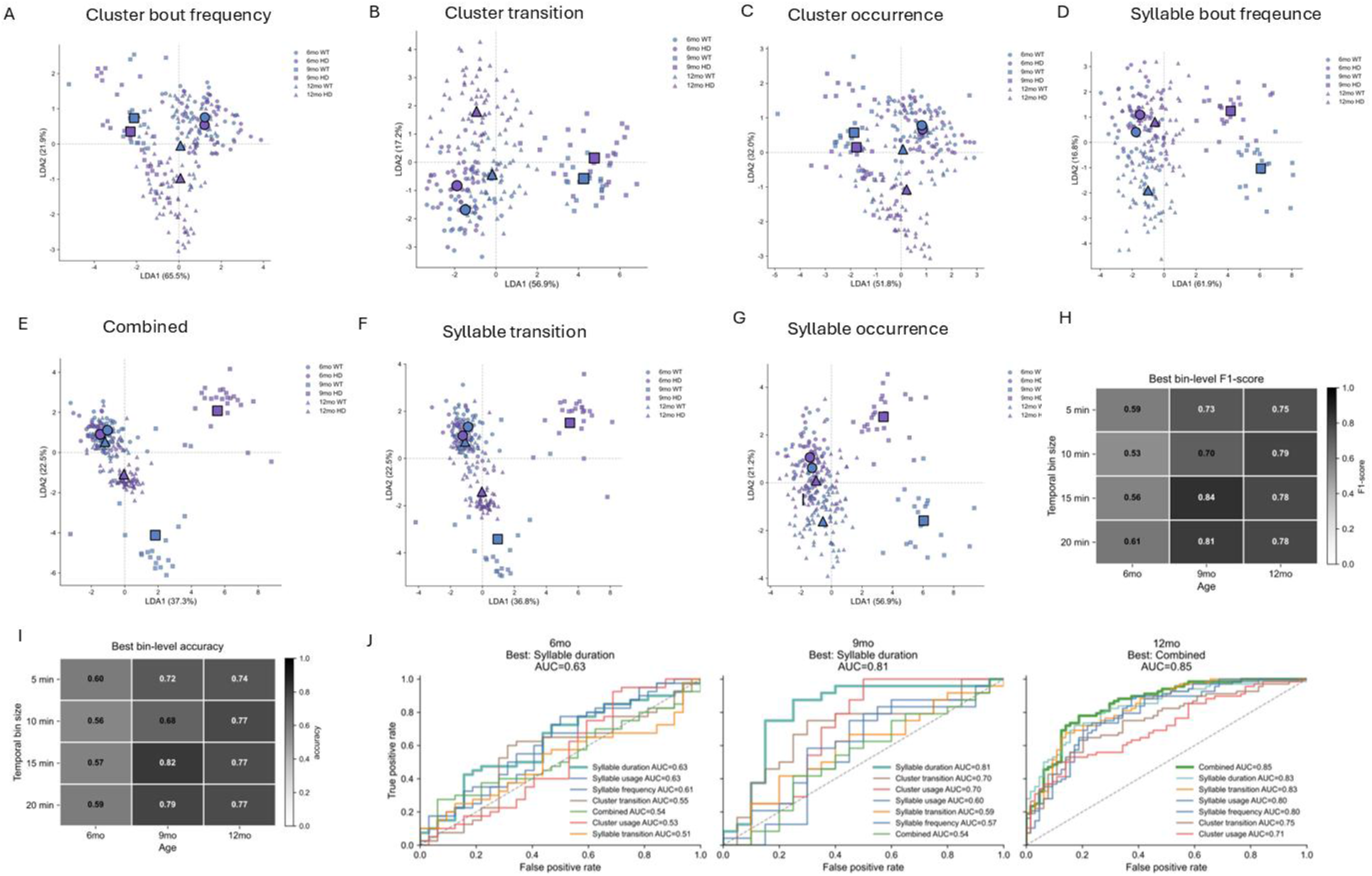
Temporal-bin optimization and feature-space screening for HD behavioral classification. **(A–G)** Linear discriminant analysis (LDA) visualization of WT and HD separation across 6-, 9-, and 12-month age groups using different behavioral feature blocks: cluster bout frequency, cluster transition, cluster occurrence, syllable bout frequency, combined features, syllable transition, and syllable occurrence. LDA was used for visualization of genotype-dependent feature-space separation. **(H, I)** Heatmaps showing the best bin-level F1 score **(H)** and classification accuracy **(I)** across temporal bin sizes and age groups. WT-versus-HD classification was performed using ElasticNet-regularized logistic regression with GroupKFold cross-validation grouped by mouse identity. **(J)** Receiver operating characteristic (ROC) curves comparing the performance of behavioral feature blocks at 6, 9, and 12 months. Syllable mean duration showed the highest classification performance at 6 and 9 months, whereas the combined feature space performed best at 12 months.

**Figure S4.**
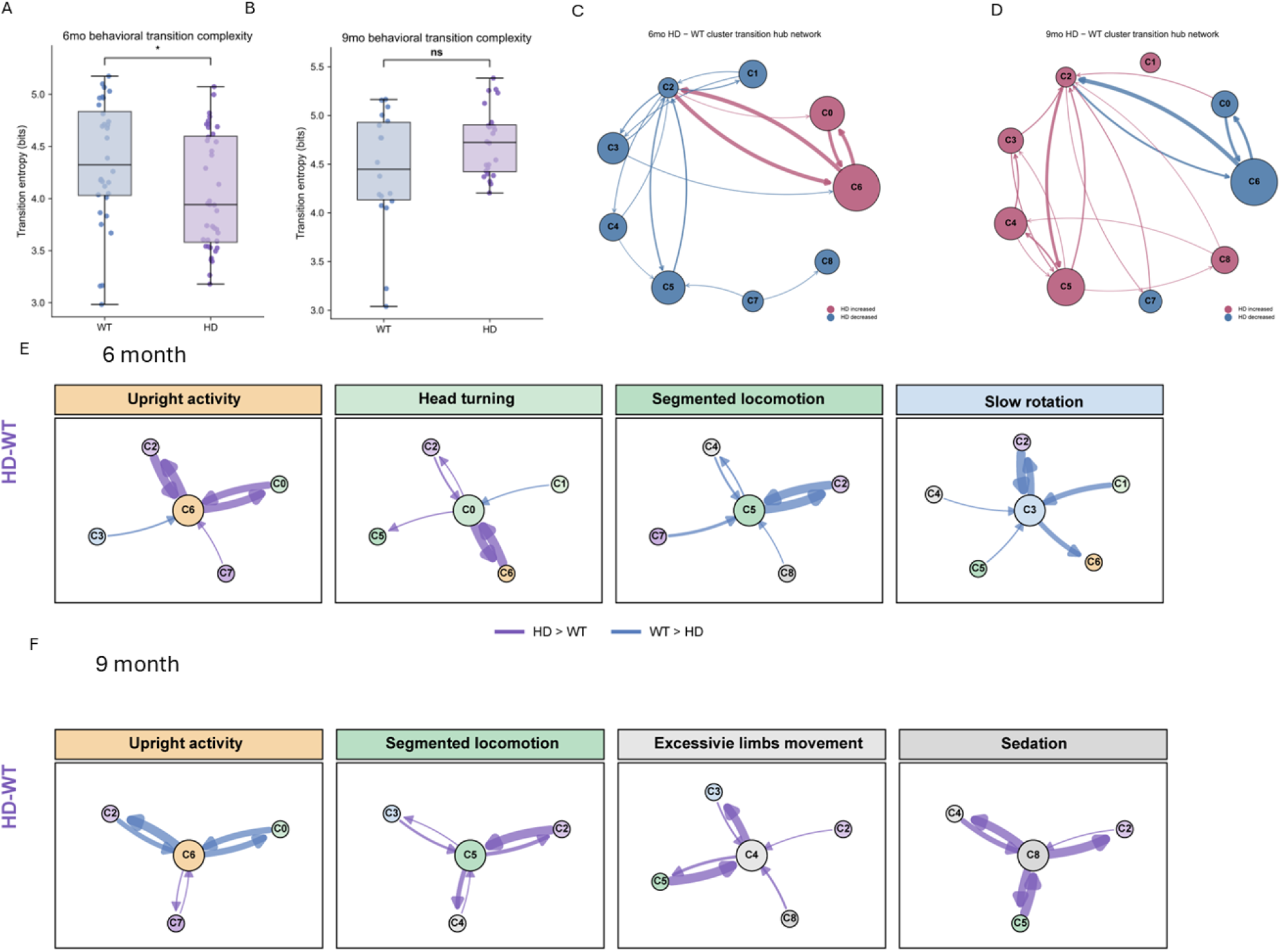
Transition-network alterations before late-stage HD behavioral disorganization. **(A, B)** Behavioral transition complexity in 6-month-old **(A)** and 9-month-old **(B)** WT and HD mice. **(C, D)** HD-minus-WT cluster transition hub networks at 6 months **(C)** and 9 months **(D)**. Node size reflects hub strength, and edge width indicates the magnitude of the HD–WT transition probability difference. **(E, F)** Connection properties of representative behavioral hubs at 6 months **(E)** and 9 months **(F)**. Purple edges indicate transitions increased in HD mice, whereas blue edges indicate transitions decreased in HD mice. Statistical significance was assessed by multiple t-tests. ns, not significant; *P < 0.05.

**Figure S5.**
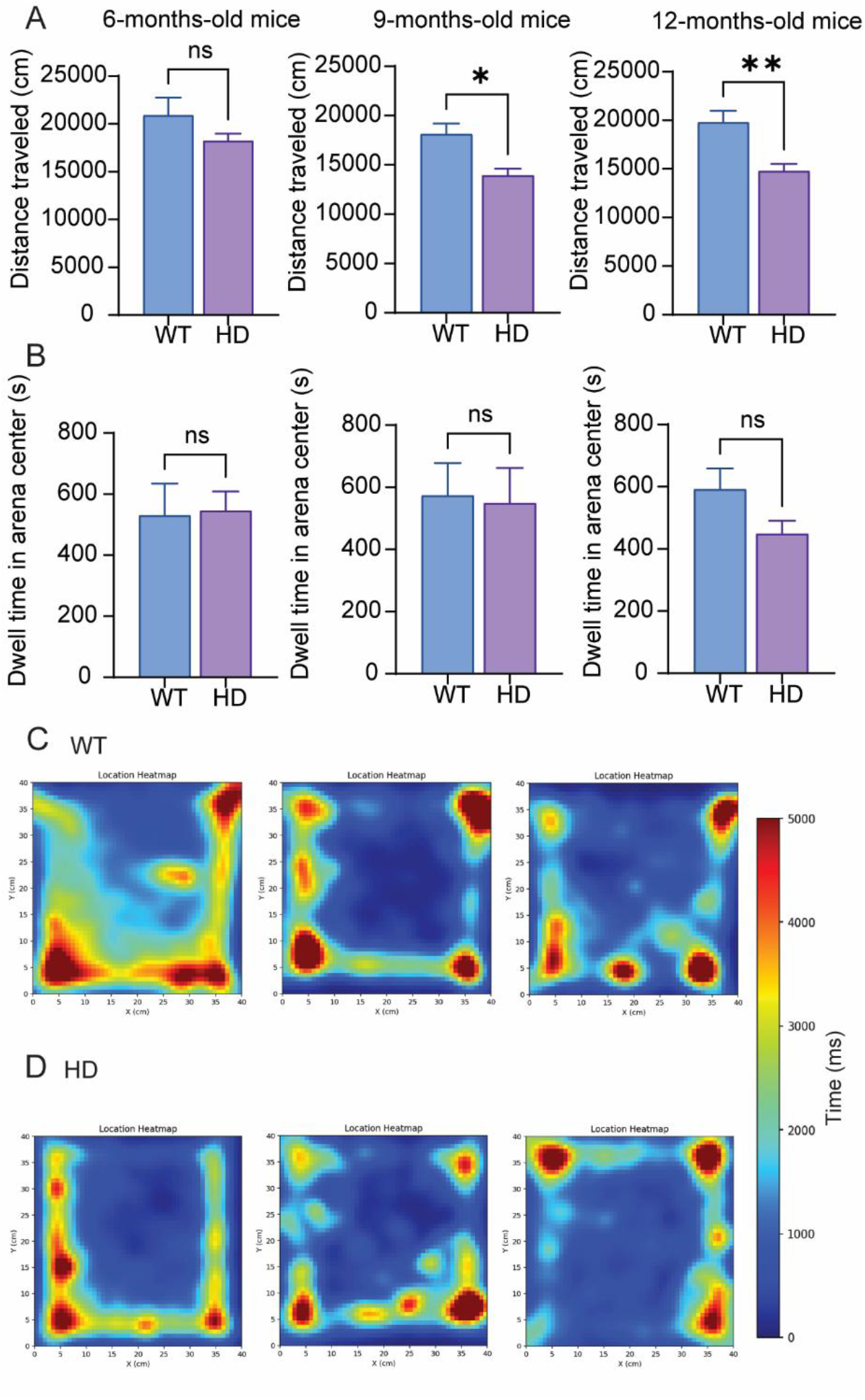
Traditional open-field measures across disease stages in WT and HD mice. **(A)** Total distance travelled during the 1-h open-field session in 6-, 9-, and 12-month-old WT and HD mice. HD mice showed reduced locomotor activity at 9 and 12 months, whereas no significant genotype difference was detected at 6 months. **(B)** Center dwell time in the open-field arena at 6, 9, and 12 months. No significant genotype differences were detected at any age. **(C, D)** Representative spatial occupancy heatmaps from individual WT **(C)** and HD **(D)** mice at 6, 9, and 12 months, shown from left to right. The color scale indicates cumulative occupancy time in milliseconds. Data in (A, B) are presented as mean ± SEM. Statistical significance was assessed by t-tests between WT and HD mice at each age. *P < 0.05, **P < 0.01; ns, not significant.

**Figure S6.**
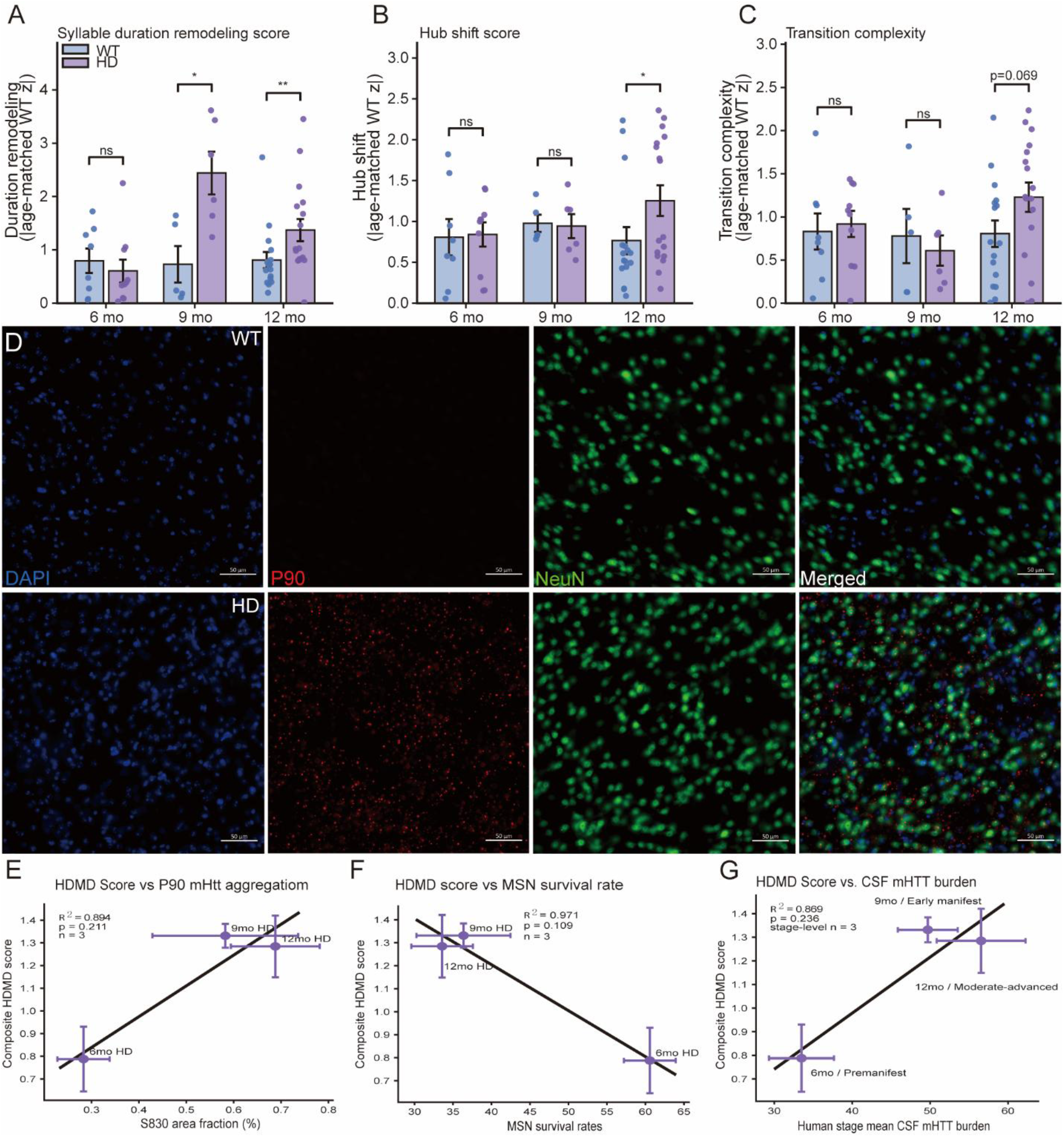
Component-level contributions and exploratory biological associations of the HD motor dysfunction score. **(A–C)** Age-dependent changes in the three absolute age-matched WT z-score components of the HDMD score: (A) syllable-duration remodeling, (B) hub shift, and (C) transition complexity. Bars show mean ± SEM; dots represent individual mice. WT and HD groups were compared using two-sided Mann–Whitney tests. *P < 0.05, **P < 0.01; ns, not significant. **(D)** Representative striatal DAPI, P90, NeuN, and merged fluorescence images from WT and HD mice. Scale bars, 50 μm. **(E–G)** Exploratory stage-level associations of the HDMD score with (E) striatal mHTT aggregate area, (F) medium spiny neuron survival, and (G) human CSF mHTT burden. Points represent stage-level means with error bars; linear fits are shown descriptively. Analyses were based on three stage-level observations and were not used for inferential validation.

**Videos S1 and S2. Representative fragmented locomotion behaviors in WT and zQ175DN HD mice.** Representative behavioral epochs assigned to the fragmented locomotion cluster in WT **(Video S1)** and zQ175DN HD **(Video S2)** mice. These videos are provided to illustrate the pose-defined behavioral pattern corresponding to this cluster; quantitative genotype differences are shown in Figure 2B.

**Videos S3 and S4. Representative slow rotation behaviors in WT and zQ175DN HD mice.** Representative behavioral epochs assigned to the slow rotation cluster in WT **(Video S3)** and zQ175DN HD **(Video S4)** mice. These videos are provided to illustrate the pose-defined behavioral pattern corresponding to this cluster; quantitative genotype differences are shown in Figure 2C.

## References

1. MacDonald ME., Ambrose CM., Duyao MP., Myers RH., Lin C., Srinidhi L., et al. A novel gene containing a trinucleotide repeat that is expanded and unstable on Huntington’s disease chromosomes. Cell 1993;72(6):971–83. Doi: 10.1016/0092-8674(93)90585-E.

2. Andhale R., Shrivastava D. Huntington’s Disease: A Clinical Review. Cureus 2022;14(8):e28484. Doi: 10.7759/CUREUS.28484.

3. Robbins AB., Ranum PT., Huerta-Ocampo I., Kuckyr M., Davidson BL. Temporal single-cell atlas of full-length Huntington’s disease mouse model defines stage-specific signatures of corticostriatal dysfunction. Mol Neurodegener 2026. Doi: 10.1186/S13024-026-00960-2.

4. Ehrlich ME. Huntington’s Disease and the Striatal Medium Spiny Neuron: Cell-Autonomous and Non-Cell-Autonomous Mechanisms of Disease. Neurotherapeutics 2012;9(2):270–84. Doi: 10.1007/s13311-012-0112-2.

5. Villanueva CB., Stephensen HJT., Mokso R., Benraiss A., Sporring J., Goldman SA. Astrocytic engagement of the corticostriatal synaptic cleft is disrupted in a mouse model of Huntington s disease. Proc Natl Acad Sci U S A 2023;120(24):e2210719120. Doi: 10.1073/PNAS.2210719120;WGROUP:STRING:PUBLICATION.

6. Menalled LB., Kudwa AE., Miller S., Fitzpatrick J., Watson-Johnson J., Keating N., et al. Comprehensive Behavioral and Molecular Characterization of a New Knock-In Mouse Model of Huntington’s Disease: ZQ175. PLoS One 2012;7(12). Doi: 10.1371/journal.pone.0049838.

7. Liu H., Zhang C., Xu J., Jin J., Cheng L., Miao X., et al. Huntingtin silencing delays onset and slows progression of Huntington’s disease: A biomarker study. Brain 2021;144(10):3101–13. Doi: 10.1093/brain/awab190.

8. Shaha R., Godad A. Huntingtin protein in health and Huntington’s disease: Molecular mechanisms, pathology and therapeutic strategies. Ageing Res Rev 2026;114. Doi: 10.1016/j.arr.2025.102984.

9. Peng Q., Wu B., Jiang M., Jin J., Hou Z., Zheng J., et al. Characterization of behavioral, neuropathological, brain metabolic and key molecular changes in zQ175 knock-in mouse model of huntington’s disease. PLoS One 2016;11(2). Doi: 10.1371/journal.pone.0148839.

10. Menalled L., El-Khodor BF., Patry M., Suárez-Fariñas M., Orenstein SJ., Zahasky B., et al. Systematic behavioral evaluation of Huntington’s disease transgenic and knock-in mouse models. Neurobiol Dis 2009;35(3):319–36. Doi: 10.1016/j.nbd.2009.05.007.

11. Koch ET., Cheng J., Ramandi D., Sepers MD., Hsu A., Fong T., et al. Deep behavioural phenotyping of the Q175 Huntington disease mouse model: effects of age, sex, and weight. BMC Biol 2024;22(1):121. Doi: 10.1186/S12915-024-01919-9.

12. Li SH., Colson TLL., Chen J., Abd-Elrahman KS., Ferguson SSG. Comparison of Huntington’s disease phenotype progression in male and female heterozygous FDNQ175 mice. Mol Brain 2023;16(1):67. Doi: 10.1186/S13041-023-01054-6.

13. Southwell AL., Smith-Dijak A., Kay C., Sepers M., Villanueva EB., Parsons MP., et al. An enhanced Q175 knock-in mouse model of Huntington disease with higher mutant huntingtin levels and accelerated disease phenotypes. Hum Mol Genet 2016;25(17):3654. Doi: 10.1093/HMG/DDW212.

14. McLean FH., Monteiro O., Lelos MJ., Ekkunagul T., Spicer RM., Rybnicek J., et al. Development of cognitive, motor, metabolic, and mutant huntingtin aggregation in the zQ175 mouse model of Huntington’s disease. Scientific Reports 2025 15:1 2025;15(1):34563-. Doi: 10.1038/s41598-025-17956-5.

15. Mathis A., Mamidanna P., Cury KM., Abe T., Murthy VN., Mathis MW., et al. DeepLabCut: markerless pose estimation of user-defined body parts with deep learning. Nat Neurosci 2018;21(9):1281–9. Doi: 10.1038/s41593-018-0209-y.

16. Pereira TD., Tabris N., Matsliah A., Turner DM., Li J., Ravindranath S., et al. SLEAP: A deep learning system for multi-animal pose tracking. Nat Methods 2022;19(4):486–95. Doi: 10.1038/s41592-022-01426-1.

17. Kabra M., Robie AA., Rivera-Alba M., Branson S., Branson K. JAABA: Interactive machine learning for automatic annotation of animal behavior. Nat Methods 2013;10(1):64–7. Doi: 10.1038/nmeth.2281.

18. Hsu AI., Yttri EA. B-SOiD, an open-source unsupervised algorithm for identification and fast prediction of behaviors. Nat Commun 2021;12(1):1–13. Doi: 10.1038/s41467-021-25420-x.

19. Luxem K., Mocellin P., Fuhrmann F., Kürsch J., Miller SR., Palop JJ., et al. Identifying behavioral structure from deep variational embeddings of animal motion. Commun Biol 2022;5(1):1267. Doi: 10.1038/S42003-022-04080-7.

20. Weinreb C., Pearl JE., Lin S., Osman MAM., Zhang L., Annapragada S., et al. Keypoint-MoSeq: parsing behavior by linking point tracking to pose dynamics. Nat Methods 2024;21(7):1329–39. Doi: 10.1038/s41592-024-02318-2.

21. Lin S., Gillis WF., Weinreb C., Zeine A., Jones SC., Robinson EM., et al. Characterizing the structure of mouse behavior using Motion Sequencing. Nat Protoc 2024;19(11):3242– 91. Doi: 10.1038/S41596-024-01015-W.

22. Markowitz JE., Gillis WF., Beron CC., Neufeld SQ., Robertson K., Bhagat ND., et al. The Striatum Organizes 3D Behavior via Moment-to-Moment Action Selection. Cell 2018;174(1):44–58.e17. Doi: 10.1016/j.cell.2018.04.019.

23. Zhang QX., Zhang YX., Jiang CC., Cheng JW., Tao SJ., Pan YL., et al. Temporal assessment of behavior in Parkinson’s visual hallucinations via a multidimensional analysis strategy. Signal Transduction and Targeted Therapy 2026 11:1 2026;11(1):146-. Doi: 10.1038/s41392-026-02651-2.

24. Oh H., Choi S., Lee J., Lee H., Shin J., Son S., et al. AI-driven decoding of naturalistic behaviors enables tailored detection of depressive-like behavior in mice. Nat Commun 2025:1–19. Doi: 10.1038/s41467-025-67559-x.

25. Zhou S., Wang X., Wang YT. 3D-AI mouse behavior analysis system has the capability to detect abnormalities in R6/1 model mice with Huntington’s disease during the pre-symptomatic phase. Front Psychiatry 2026;17:1749543. Doi: 10.3389/FPSYT.2026.1749543/TEXT.

26. Ye S., Filippova A., Lauer J., Schneider S., Vidal M., Qiu T., et al. SuperAnimal pretrained pose estimation models for behavioral analysis. Nature Communications 2024 15:1 2024;15(1):5165-. Doi: 10.1038/s41467-024-48792-2.

27. Mathis A., Mamidanna P., Cury KM., Abe T., Murthy VN., Mathis MW., et al. DeepLabCut: markerless pose estimation of user-defined body parts with deep learning. Nat Neurosci 2018;21(9):1281–9. Doi: 10.1038/s41593-018-0209-y.

28. Weinreb C., Pearl JE., Lin S., Osman MAM., Zhang L., Annapragada S., et al. Keypoint-MoSeq: parsing behavior by linking point tracking to pose dynamics. Nat Methods 2024;21(7):1329–39. Doi: 10.1038/s41592-024-02318-2.

29. Huang K., Han Y., Chen K., Pan H., Zhao G., Yi W., et al. A hierarchical 3D-motion learning framework for animal spontaneous behavior mapping. Nature Communications 2021 12:1 2021;12(1):2784-. Doi: 10.1038/s41467-021-22970-y.

30. Wu Q., Yao M., Liu H., Kakazu A., Ouyang Y., Liu C., et al. Progressively reduced cerebral oxygen metabolism and elevated plasma NfL levels in the zQ175DN mouse model of Huntington’s disease. Exp Neurol 2025;394. Doi: 10.1016/j.expneurol.2025.115461.

31. Sebastianutto I., Cenci MA., Fieblinger T. Alterations of striatal indirect pathway neurons precede motor deficits in two mouse models of Huntington’s disease. Neurobiol Dis 2017;105:117–31. Doi: 10.1016/J.NBD.2017.05.011.

32. Weinreb C., Kannan LT., Newman-Boulle A., Sainburg T., Gillis WF., Plotnikoff A., et al. Spontaneous behavior is a succession of self-directed tasks. Neuron 2026;114(5):922. Doi: 10.1016/J.NEURON.2025.11.021.

33. Jørgensen SH., Ejdrup AL., Lycas MD., Posselt LP., Madsen KL., Tian L., et al. Behavioral encoding across timescales by region-specific dopamine dynamics. Proc Natl Acad Sci U S A 2023;120(7):e2215230120. Doi: 10.1073/PNAS.2215230120;ISSUE:ISSUE:DOI.

34. Zhang B., Geddes CE., Jin X. Complementary corticostriatal circuits orchestrate action repetition and switching. Science Advances 2025;11(21). Doi: 10.1126/SCIADV.ADT0854;PAGEGROUP:STRING:PUBLICATION.

35. Cruz BF., Guiomar G., Soares S., Motiwala A., Machens CK., Paton JJ. Action suppression reveals opponent parallel control via striatal circuits. Nature 2022 607:7919 2022;607(7919):521–6. Doi: 10.1038/s41586-022-04894-9.

36. Ehrnhoefer DE., Wong BKY., Hayden MR. Convergent pathogenic pathways in Alzheimer’s and Huntington’s diseases: shared targets for drug development. Nature Reviews Drug Discovery 2011 10:11 2011;10(11):853–67. Doi: 10.1038/nrd3556.

37. Korchynska S., Rebernik P., Pende M., Boi L., Alpár A., Tasan R., et al. A hypothalamic dopamine locus for psychostimulant-induced hyperlocomotion in mice. Nature Communications 2022 13:1 2022;13(1):5944-. Doi: 10.1038/s41467-022-33584-3.

38. Ibrahim KS., Mestikawy S El., Abd-Elrahman KS., Ferguson SSG. VGLUT3 Deletion Rescues Motor Deficits and Neuronal Loss in the zQ175 Mouse Model of Huntington’s Disease. Journal of Neuroscience 2023;43(23):4365–77. Doi: 10.1523/JNEUROSCI.0014-23.2023.

39. Scannell JW., Bosley J., Hickman JA., Dawson GR., Truebel H., Ferreira GS., et al. Predictive validity in drug discovery: what it is, why it matters and how to improve it. Nature Reviews Drug Discovery 2022 21:12 2022;21(12):915–31. Doi: 10.1038/s41573-022-00552-x.

40. von Ziegler LM., Roessler FK., Sturman O., Waag R., Privitera M., Duss SN., et al. Analysis of behavioral flow resolves latent phenotypes. Nature Methods 2024 21:12 2024;21(12):2376–87. Doi: 10.1038/s41592-024-02500-6.

41. Whiteway MR., Biderman D., Friedman Y., Dipoppa M., Buchanan EK., Wu A., et al. Partitioning variability in animal behavioral videos using semi-supervised variational autoencoders. PLoS Comput Biol 2021;17(9):e1009439. Doi: 10.1371/JOURNAL.PCBI.1009439.

42. Vollert J., Macleod M., Dirnagl U., Kas MJ., Michel MC., Potschka H., et al. The EQIPD framework for rigor in the design, conduct, analysis and documentation of animal experiments. Nature Methods 2022 19:11 2022;19(11):1334–7. Doi: 10.1038/s41592-022-01615-y.

43. Estevez-Fraga C., Scahill RI., Durr A., Leavitt BR., Roos RAC., Langbehn DR., et al. Composite UHDRS Correlates With Progression of Imaging Biomarkers in Huntington’s Disease. Movement Disorders 2021;36(5):1259–64. Doi: 10.1002/MDS.28489;WGROUP:STRING:PUBLICATION.

44. Rodrigues FB., Byrne LM., Tortelli R., Johnson EB., Wijeratne PA., Arridge M., et al. Mutant huntingtin and neurofilament light have distinct longitudinal dynamics in Huntington’s disease. Sci Transl Med 2020;12(574):2888. Doi: 10.1126/scitranslmed.abc2888.

